# Out of the Qinghai-Tibetan Plateau and get flourishing - the evolution of *Neodon* voles (Rodentia: Cricetidae) revealed by systematic sampling and low coverage whole genome sequencing

**DOI:** 10.1101/2019.12.12.873679

**Authors:** Shaoying Liu, Chengran Zhou, Tao Wan, Guanliang Meng, W. Robert W. Murphy, Zhengxin Fan, Mingkun Tang, Yang Liu, Tao Zeng, Shunde Chen, Yun Zhao, Shanlin Liu

## Abstract

*Neodon*, genus of a short time evolutionary history, was reported to be diverged from its relatives in early stage of Pleistocene. Only 4 species were well documented in *Neodon* for a long period of time until last years when a systematic work described and added three new species, adjusted three species used to belong to *Lasiopodomys, Phaiomys, Microtus* to *Neodon* and removed one species (*Neodon juldaschi*) to genus *Blanfordimys*, leading to a total of eight species recorded in *Neodon*. To gain a better insight into the phylogeny and ecology of *Neodon*, we have systematically sampled *Neodon* species along the whole Hengduan and Himalayan Mountains in the last 20 years. In addition to morphological identification, we generated 1x - 15x whole genome sequencing (WGS) data and achieved the mitochondrial genomes and an average of 5,382 nuclear genes for each morpho-species. Both morphology and phylogeny results supported an extra six new species in *Neodon* (nominated *Neodon shergylaensis* sp. nov., *N. namchabarwaensis* sp. nov., *N. liaoruii* sp. nov., *N. chayuensis* sp. nov., *N. bomiensis* sp. nov., and *N. bershulaensis* sp. nov.). This is the first study that included *Neodon* samples covering its entire distribution area in China and this systematic sampling also revealed a long-time underestimation of *Neodon*’s diversity, and suggested its speciation events linked highly to founder event via dispersal (from Plateau to surrounding mountains). The results also revealed that the Qinghai-Tibetan Plateau is the center of origin of *Neodon*, and the impetus of speciation include climate change, isolation of rivers and mountains.

---

The Tibetan-Himalayan region (THR), comprised of the Himalayas, the Hengduan Mountains (HD) biodiversity hotspot and the Qinghai-Tibetan Plateau (QTP), consists of a series of parallel alpine ridges and deep river valleys forming dramatic ecological stratification and environmental heterogeneity (Marchese 2015; Muellner-Riehl 2019; Myers, et al. 2000). The complex topographical features and physical boundaries lead to the geographic isolation of biota that limits or ceases gene flow. This can drive speciation. The topographical complexity and vast territory also constrain fieldwork, and this affects estimates of species richness and testing hypotheses on species’ interactions, and especially for species with limited dispersal abilities. The QTP is regarded to be a “museum of evolution” and a “cradle of evolution” (Moreau and Bell 2013; Mosbrugger, et al. 2018; Tamma and Ramakrishnan 2015; Xing and Ree 2017) because of its high percentages of ancient species and biodiversity. Evidence is accumulating that the QTP is the center of origin and accumulation for many organisms with particular biogeographical relationships to other Palearctic regions, and this support the “out of Tibet” hypothesis (Jia, et al. 2012; Mosbrugger, et al. 2018; Pisano, et al. 2015; Wang, et al. 2014; Weigold 2005). Thus, the Tibetan-Himalayan region provides critical clues to how geology and climate together drive the evolution.

Voles and lemmings are one of the youngest groups of rodents and the most recent ancestor of *Neodon* and *Microtus* (Rodentia: Cricetidae) was dated at about 7 million years ago (Mya) (Abramson, et al. 2009; Lv, et al. 2016). Speciation within *Neodon* was driven by orogenesis in the HD region from the late Miocene, when mountains surrounding QTP have reached its current elevations (~ 8 Mya), to the late Pliocene (~2 Mya) (Mosbrugger, et al. 2018; Muellner-Riehl 2019; Xing and Ree 2017). *Neodon* was erected by Horsfield in 1841 with only four well-documented species (*N. sikimensis, N. irene, N. forresti* and *N. juldaschi*). The genus occurs only in the Himalayas, HD and QTP (Liu, et al. 2017). A long-running taxonomic debate involved *Neodon*’s phylogenetic position—either as a subgenus of *Microtus* (Allen 1940; Carleton and Musser 1995; Gromov and Polyakov 1977) or as a subgenus of *Pitymys* (Corbet 1978; Ellerman 1949; Ellerman and Morrison-Scott 1951). However, recent morphological and molecular evidence confirmed *Neodon* to be a monophyletic genus (Ellerman 1941; Liu, et al. 2017; Liu, et al. 2012; Musser and Carleton 2005), and having far more than four species. For example, all but two species in Arvicolinae (Cricetidae) on the QTP and surrounding high elevation areas belong to *Neodon*.

To gain better insights into the phylogenetic status and diversity of *Neodon*, as well as the role played by the QTP in driving biogeography and diversification, we report on a collection of specimens of small mammals taken from throughout the distribution *Neodon*, and especially in the Himalayas, over the past 20 years. Sampling, which covers tens of thousands of square kilometers, achieved more than 2,000 samples, of which 193 samples were morphologically identified to be *Neodon* (Fig. 1, Supplementary Appendix S1). In addition to morphological and geographic data, we provide 1X–15X whole genome sequencing (WGS) data for each representative morphological species (Fig. 2, Supplementary Table S1), identify and describe six new species, demonstrate the underestimation of diversity in China and reveal that drastic climate change and topography of THR, as well as the founder event via dispersal, influence the diversification. These results suggest that similar investigations of other small mammals will discover greater diversity and lead to identifying the common driver(s) of patterns and processes in the THR ecosystem.

**Figure 1.**
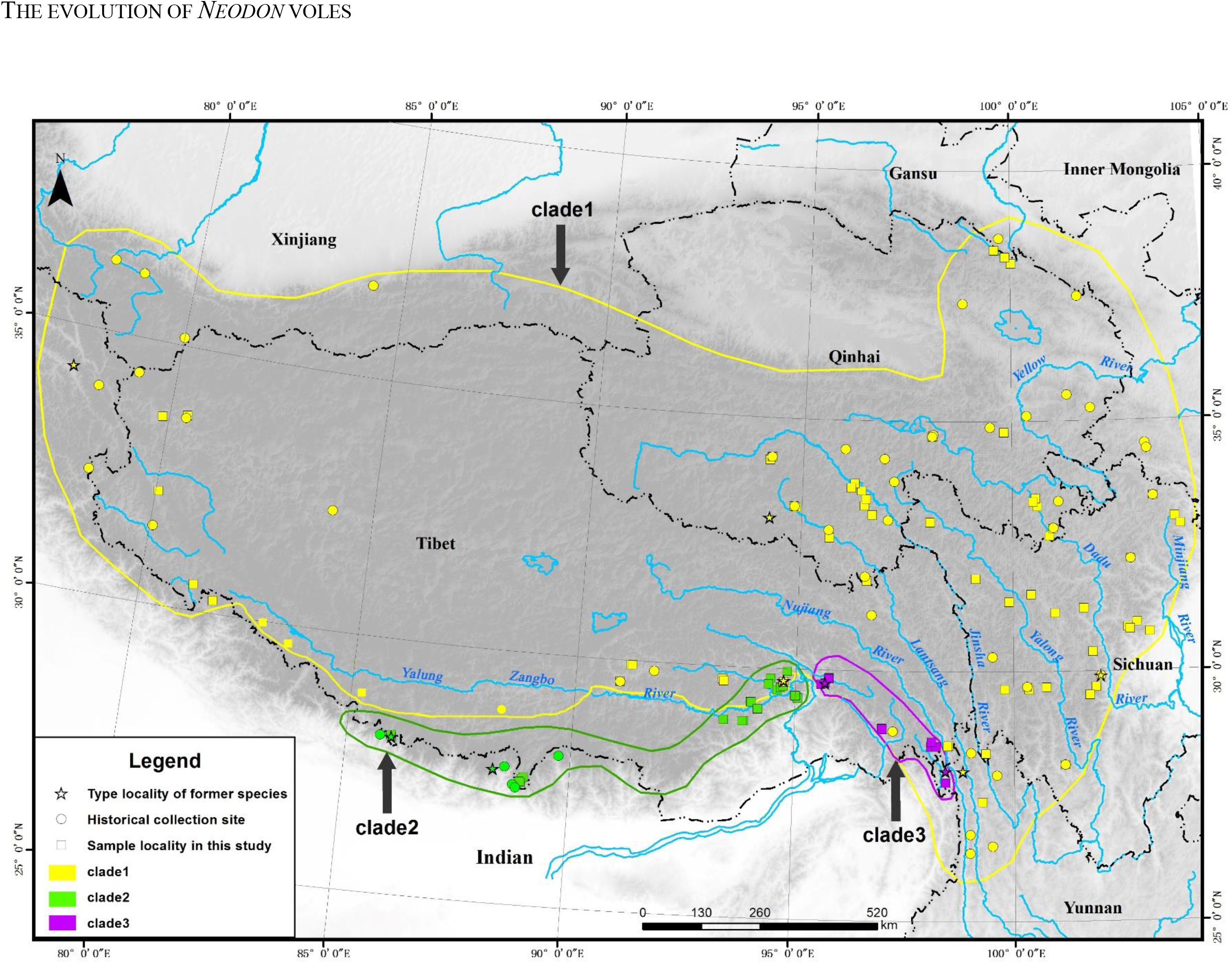
Geographical distribution of *Neodon* in this study. Approximate extent of occurrence of each clade is shown as colored lines (clade 1: yellow; clade 2: green, clade 3: purple, refer to Fig. 4 for clade information). Stars show the type localities of former species. Circles show the historical collection sites. Squares show the distribution of newly collected specimens.

**Figure 2.**
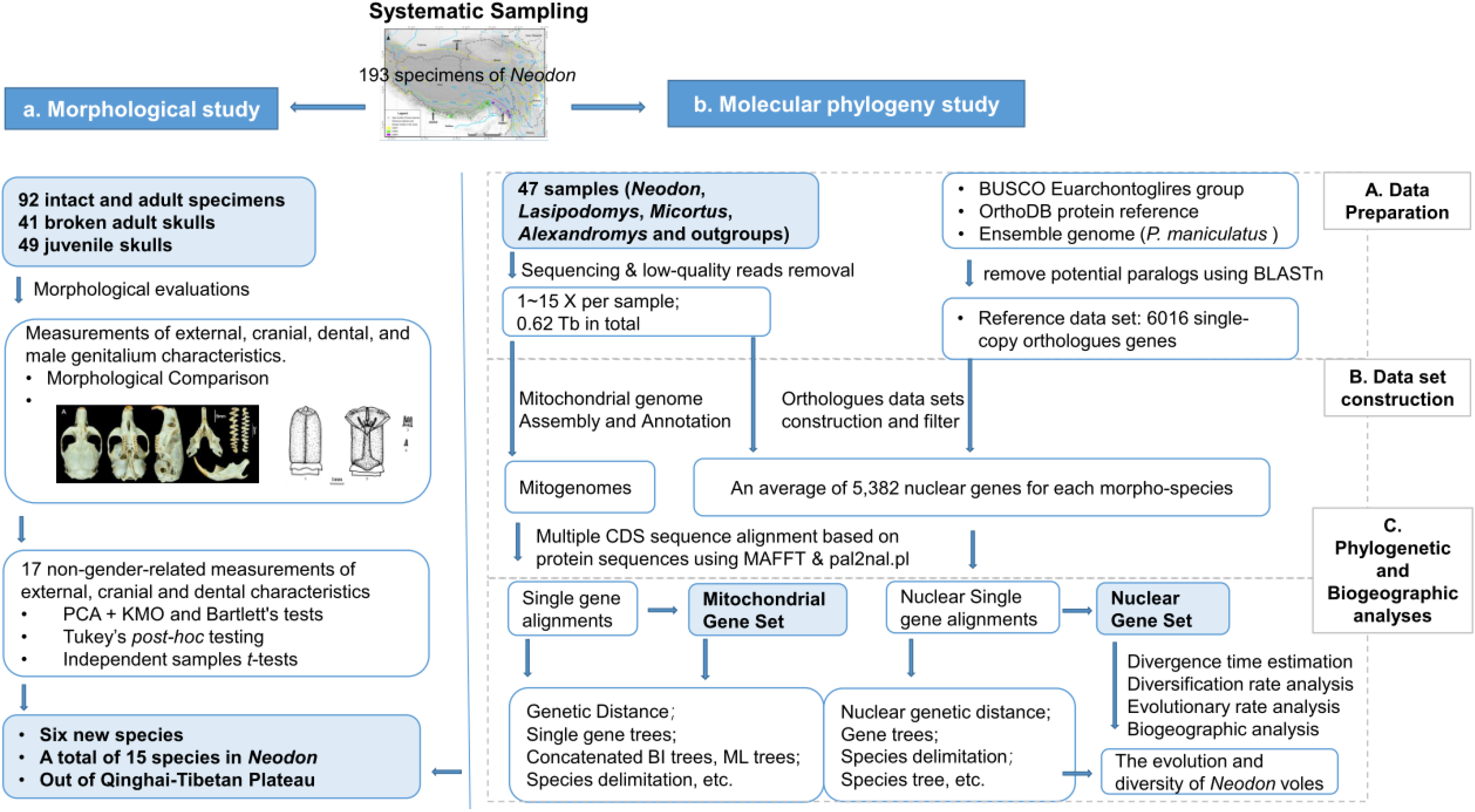
Schematic pipeline illustrating the workflow of morphological analysis and molecular phylogeny analysis after systematic sampling. a) Workflow of morphological analysis. b) Workflow of molecular analysis, including: A. sample preparation for sequencing and reference data set construction; B. mitochondrial and nuclear data set construction; and C. phylogenetic analysis, divergence time estimation, diversification analysis, incomplete lineage sorting (ILS) testing, species delimitation, et al. The combined results of both morphological analysis and molecular phylogeny analysis shows that *Neodon* has 15 species, including six new species identified herein.

## Materials and Methods

### Ethics statement

All samples were obtained following Guidelines of the American Society of Mammalogists (ASM guidelines) (Sikes and Gannon 2011) and the laws and regulations of China for the implementation of the protection of terrestrial wild animals (State Council Decree 1992). Collecting protocols were approved by the Ethics Committee of the Sichuan Academy of Forestry (no specific permit number). Voucher specimens were deposited in the Sichuan Academy of Forestry, Chengdu, China.

### Sample information

We included a total of 193 specimens of *Neodon* that collected in QTP for analyses. The collection comprised 54 juveniles and 139 adults, of which 99 had intact skulls and were used for statistical analysis of morphology. To verify the morphological identification and further investigate the evolutionary history of *Neodon*, we generated WGS data for a total of 48 specimens, representing 15 putative species of *Neodon*, four species of *Lasiopodomys*, three species of *Eothenomys*, one species of *Craseomys*, one species of *Caryomys*, one species of *Myodes*, and seven species of *Microtus* (Supplementary Table S1). All specimens were housed at the Sichuan Academy of Forestry (SAF).

### Morphology analyses

Morphological evaluations for the 92 intact adult specimens identified, *N. clarkei*, *N. fuscus*, *N. irene*, *N. leucurus*, *N. linzhiensis*, *N. nyalamensus*, *N. medogensis* and *N. sikimensis*, plus one previously evaluated unnamed (*N. forresti*) and six tentatively new species. We also included an additional 41 broken adult skulls and 49 juvenile skulls of the putative six new species for comparison.

#### Measurement of morphological characteristics

For all the specimens, we collected the external, cranial, and dental characteristics of which the external measurements were recorded in the field on freshly captured specimens to an accuracy of 0.5 mm and the cranial and dental characteristics were measured using a Vernier caliper to an accuracy of 0.02 mm in lab. The measurements included previously used characteristics (Supplementary Table S2) (Liu, et al. 2017). For males, we recorded characteristics of genitalium, which provides significant clues to interrelationships of species (Hooper 1958; Hooper and Hart 1962). We prepared the glans penes following the canonical methods (Hooper 1958; Lidicker Jr 1968) and characterized bacular structures (Hooper 1958; Yang and Fang 1988; Yang, et al. 1992) with several measurements (Supplementary Table S2).

Morphometric variation of 17 non-gender-related measurements of adult specimens was analyzed using principal component analyses (PCAs) in SPSS v.17.0. We employed Kaiser-Meyer-Olkin (KMO) and Bartlett’s tests to check the fitness of the PCA analysis followed by Tukey’s test and independent samples *t*-tests to assess statistical differences.

### High throughput sequencing (HTS)

We extracted genomic DNA for each specimen from muscle tissues using Gentra Puregene Tissus Kit (Qiagen, Valencia, CA) according to the manufacturer’s protocol, and then generated > 10 Gb data for one representative specimen of each morphological species and about 3 Gb data (1X) for the other specimens from the same species. Low-quality reads of high-coverage specimens were removed if they met one or more following criteria: 1) an N-content of more than 10%; 2) adapter contaminated reads (reads overlapping more than 50% with the adapter sequence, with a maximum of 1 bp mismatches to the adaptor sequence); and 3) more than 20% of the read-length below Q10. The criteria of low-quality read-filtering of other specimens was put in Supplementary Table S1.

#### Mitochondrial genome assembly and annotation

We assembled and annotated the mitogenome for each sample using MitoZ (Meng, et al. 2019) with about 3 Gb (~1X) clean data. For *Neodon* and other closely-related genera, we also downloaded available cytochrome b (*cytb*) and cytochrome c oxidase subunit I (*cox1*) genes from GenBank (accessed in Oct. 2018), the two most widely used genetic marker for small mammals (Jia, et al. 2012; Liu, et al. 2017; Lv, et al. 2018; Zhang, et al. 2016).

#### Construction of orthologues data sets

To obtain orthologous nuclear genes for each sample (Fig. 2), we at first obtained 6,192 lineage-specific single-copy orthologs for the euarchontoglires group by using the Benchmarking Universal Single-Copy Orthologs (BUSCO) database (v9) (Simão, et al. 2015). Then, we downloaded the genome of the North American deer mouse (Creicetidae, *Peromyscus maniculatus*), the most closely related species of *Neodon* available in the BUSCO database, from Ensemble (Hubbard, et al. 2002) to obtain its corresponding full gene regions (including exons and introns). Next, genes with high similarity homologs (BLASTn v2.6.0+ with e value < 1e-5) (Altschul, et al. 1990) within the current gene set were removed to avoid mapping uncertainty in the next step. Next, we aligned WGS data of each sample to the *Peromyscus maniculatus* genome using BWA-MEM (Li 2013) with default parameters and obtained their corresponding genes using consensus calling function in bcftools v.1.8 (Li, et al. 2009) (detailed in Supplementary Appendix S1). Finally, high quality CDS—no internal stop codons, ‘N’ content smaller than 20% and present in more than 50% of the total samples—were filtered out for subsequent analyses. We firstly aligned their corresponding protein sequences using MAFFT v.7.313 (Katoh and Standley 2013) and then achieved the CDS alignments using PAL2NAL (Suyama, et al. 2006) based on the protein alignments. Final alignments were given in Supplementary Appendices S2-S5, available on Dryad.

#### Genetic distance calculation

We calculated the Kimura 2-parameter genetic distances of each nuclear and mitochondrial gene using the dist.dna function in the R ape v.1.1-1 package (Paradis, et al. 2004). The average genetic distance of each taxonomic group (e.g. within species, within genus) was calculated.

#### Species Delimitation

In addition to morphological identification, we conducted species delimitation using two sequence-based methods: the clustering-based Bayesian implementation of the Poisson Tree Processes (bPTP) analysis (Zhang, et al. 2013) using both mitochondrial and nuclear trees and the similarity-based Automatic Barcode Gap Discovery (ABGD) analysis using mitochondrial data (Puillandre, et al. 2012).

#### Phylogenetic inference

We inferred the phylogeny using RAxML (Stamatakis 2014) with GTR+GAMMA+I model and 100 bootstrap replicates for each gene. Next, the final species tree was achieved using ASTRAL-III (Zhang, et al. 2018) based on the multispecies coalescent model and the bootstrap support of each node was estimated by the multi-locus resampling method. SVDquartets (parameters of “eval Quartets=1e+6 bootstrap=standard”) implemented in PAUP v.4.0a164 (Chifman and Kubatko 2014; Swofford 2001) was also utilized to estimate the species tree with the same dataset to validate the results. Simultaneously, we concatenated the CDS alignments to generate a “supergene” alignment for each species and used MrBayes v.3 (Huelsenbeck and Ronquist 2001) and RAxML to construct concatenated trees. Alignments included a “mitochondrial Gene Set” and “nuclear Gene Set”. Branch lengths of final species tree were re-estimated in units of substitutions per site by constraining alignments to the species tree topology using ExaML v.3.0.21 (Kozlov, et al. 2015). Trees were outgroup-rooted with species in *Eothenomys, Craseomys*, *Caryomys* and *Myodes*.

The divergence time of the species tree were estimated based on the second codon sites of the nuclear genes using the Bayesian relaxed clock method MCMCTree implemented in the PAML v.4.9h package (Yang 2007) with the approximate likelihood calculation of ‘REV’ (GTR, model=7) model and with fossil calibration points taken from records in the Paleobiology Database (Available: https://paleobiodb.org, Accessed 2018 Dec 12) and the timetree database (Kumar, et al. 2017) (detailed in Supplementary Appendix S1 and Table S3).

### Evolutionary and biogeographic analyses

#### Incomplete lineage sorting investigation

Because incomplete lineage sorting (ILS) may cause incongruence in phylogenetic trees inferred from the different datasets, we scanned for the presence of ILS spanning the evolution of *Neodon* with the nuclear data set of all 28 taxa (including outgroups) using DiscoVista (Sayyari, et al. 2018). The correlation between ILS content and the inner-node branch length were calculated based on the linear models and Pearson test in R (Field, et al. 2012) and visualized using ggplot2 (Wickham 2016).

#### Evolutionary rate analysis

To investigate the evolutionary rate of different clades within *Neodon* and the relationship to their corresponding living conditions, we calculated the evolutionary rates of mitochondrial genes and a nuclear gene set including 100 genes with top ASTRAL gene tree scores using the “several ω ratio” branch model (model = 2) implemented in the PAML package (v.4.9h) (Yang 2007) with the external and internal branches being set as foreground and background, respectively. The evolutionary rate of each branch was visualized using R package ggtree (Yu, et al. 2017).

#### Diversification rate analysis

To assess the diversification of *Neodon* through time, we generated Log-Lineage through time (LTT) plots for the time-calibrated phylogeny (non-*Neodon* species were pruned), as well as for 100 simulated trees of the same age and taxon richness, using Phytools (Revell 2012). For generating simulated trees, the ‘Yule’ (pure-birth) model and birth-death model were compared using Akaike information criterion (AIC). Next, 100 simulated trees were generated using ‘pbtree’ implemented in the Phytools under a Yule model and a constant rate model with the speciation rate of 0.487955 estimated using ‘fit.bd’.

#### Biogeographic analysis

Extended outgroups were pruned from the tree so that only *Neodon*, *Alexandormys*, *Microtus* and *Lasiopodomys* were analyzed. We used BioGeoBears v.1.1.2 (Matzke 2013a) for biogeographic reconstruction based on the species tree and assigned the species to one or two of the following biogeographical regions according to their distributions with the Tsangpo River and the Mekong-Salween rivers divide being used as the border for EH-H and H-HD, respectively: P (the QTP); EH (Eastern Himalayan Mountains); H (Himalayas); HD (Hengduan Mountains); O (area out of THR). The maximum range size was set to 2 because no extant species occurs in ≥ 3 biogeographical regions as defined here. We tested a total of six models of biogeographical reconstruction in the likelihood framework in BioGeoBears, including dispersal‐extinction cladogenesis model (DEC) (Ree and Smith 2008), dispersal–vicariance analysis (DIVA) (Ronquist 1997) and the BayArea model (Landis, et al. 2013), plus all three models separately under the possibility of founder events (+J) (Matzke 2014; Matzke 2013b). AIC scores was employed to compare the fit of different models.

#### Data records

Data that support our findings were published under the International Nucleotide Sequence Database Collaboration BioProject PRJNA564473 (ncbi.nlm.nih.gov/bioproject/?term=PRJNA564473) and CNGB Nucleotide Sequence Archive (CNSA) project CNP0000173 (db.cngb.org/search/project/CNP0000173).

## Results

### Morphological comparison

#### Comparisons of skull, teeth and bacular structures

Initial observations of skulls, teeth and bacular structures revealed 15 distinct patterns (Table 1, Fig. 3–4, Supplementary Fig. S1-2) each representing a putative species of *Neodon*. This included eight described species, one previously evaluated unnamed taxon and six tentatively new species. Skulls were compared in Supplementary Fig. 1 and molars and glans penes for all species of *Neodon* in Fig. 4 and Table 1. Molar patterns and morphology of glans penes clearly distinguished all unidentified species of *Neodon*.

**Table 1.**
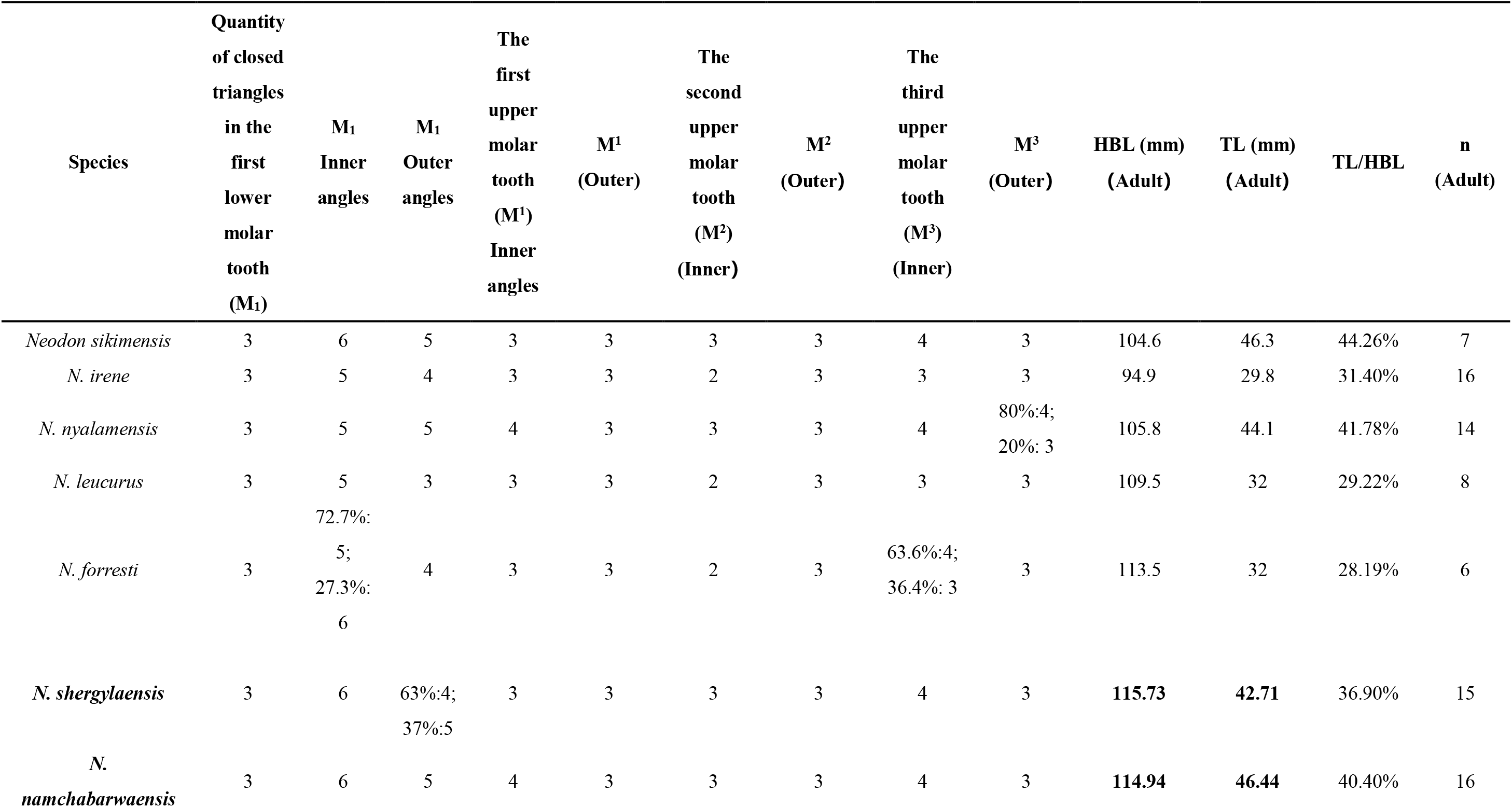
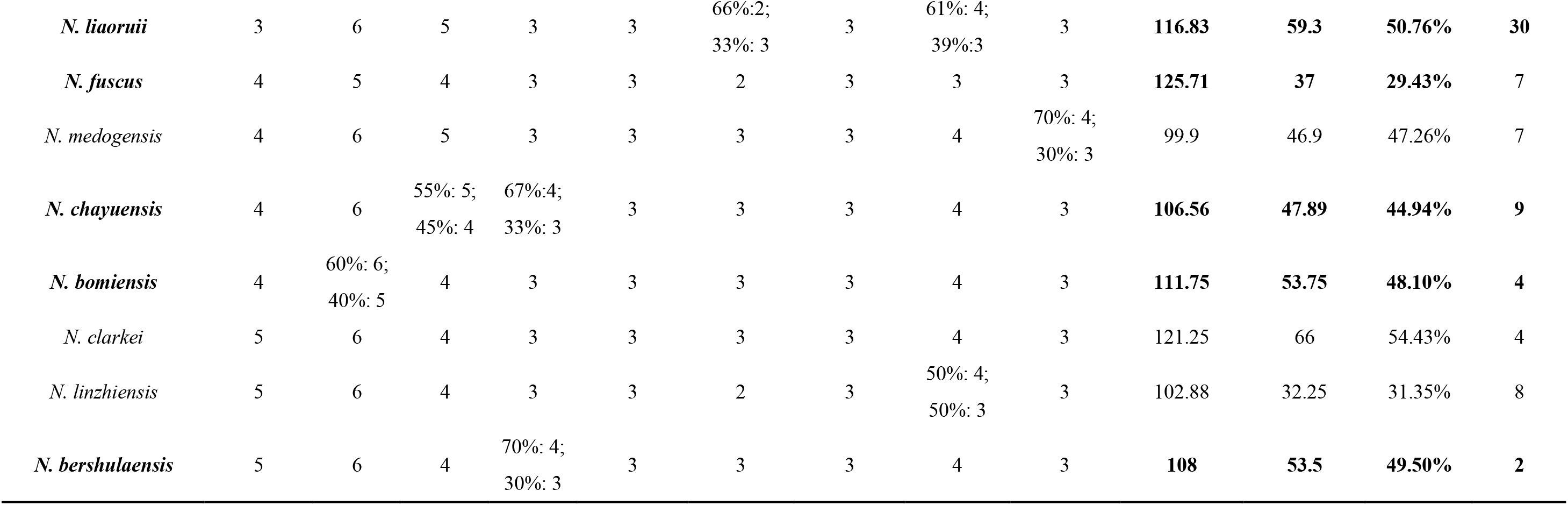
Morphology comparison of 15 species of *Neodon*

| Species | Quantity<br>of closed<br>triangles<br>in the<br>first<br>lower<br>molar<br>tooth<br>(M<sub>1</sub>) | M<sub>1</sub><br>Inner<br>angles | M<sub>1</sub><br>Outer<br>angles | The<br>first<br>upper<br>molar<br>tooth<br>(M<sup>1</sup>)<br>Inner<br>angles | M<sup>1</sup><br>(Outer) | The<br>second<br>upper<br>molar<br>tooth<br>(M<sup>2</sup>)<br>(Inner) | M<sup>2</sup><br>(Outer) | The<br>third<br>upper<br>molar<br>tooth<br>(M<sup>3</sup>)<br>(Inner) | M<sup>3</sup><br>(Outer) | HBL (mm)<br>(Adult) | TL (mm)<br>(Adult) | TL/HBL | n<br>(Adult) |
| --- | --- | --- | --- | --- | --- | --- | --- | --- | --- | --- | --- | --- | --- |
| *Neodon sikimensis* | 3 | 6 | 5 | 3 | 3 | 3 | 3 | 4 | 3 | 104.6 | 46.3 | 44.26% | 7 |
| *N. irene* | 3 | 5 | 4 | 3 | 3 | 2 | 3 | 3 | 3 | 94.9 | 29.8 | 31.40% | 16 |
| *N. nyalamensis* | 3 | 5 | 5 | 4 | 3 | 3 | 3 | 4 | 80%:4;<br>20%: 3 | 105.8 | 44.1 | 41.78% | 14 |
| *N. leucurus* | 3 | 5 | 3 | 3 | 3 | 2 | 3 | 3 | 3 | 109.5 | 32 | 29.22% | 8 |
| *N. forresti* | 3 | 72.7%:<br>5;<br>27.3%:<br>6 | 4 | 3 | 3 | 2 | 3 | 63.6%:4;<br>36.4%: 3 | 3 | 113.5 | 32 | 28.19% | 6 |
| *N. shergylaensis* | 3 | 6 | 63%:4;<br>37%:5 | 3 | 3 | 3 | 3 | 4 | 3 | **115.73** | **42.71** | 36.90% | 15 |
| *N.*<br>*namchabarwaensis* | 3 | 6 | 5 | 4 | 3 | 3 | 3 | 4 | 3 | **114.94** | **46.44** | 40.40% | 16 |
| *N. liaoruii* | 3 | 6 | 5 | 3 | 3 | 66%:2;<br>33%: 3 | 3 | 61%: 4;<br>39%:3 | 3 | **116.83** | **59.3** | **50.76%** | **30** |
| *N. fuscus* | 4 | 5 | 4 | 3 | 3 | 2 | 3 | 3 | 3 | **125.71** | **37** | **29.43%** | 7 |
| *N. medogensis* | 4 | 6 | 5 | 3 | 3 | 3 | 3 | 4 | 70%: 4;<br>30%: 3 | 99.9 | 46.9 | 47.26% | 7 |
| *N. chayuenensis* | 4 | 6 | 55%: 5;<br>45%: 4 | 67%:4;<br>33%: 3 | 3 | 3 | 3 | 4 | 3 | **106.56** | **47.89** | **44.94%** | **9** |
| *N. bomiensis* | 4 | 60%: 6;<br>40%: 5 | 4 | 3 | 3 | 3 | 3 | 4 | 3 | **111.75** | **53.75** | **48.10%** | **4** |
| *N. clarkei* | 5 | 6 | 4 | 3 | 3 | 3 | 3 | 4 | 3 | 121.25 | 66 | 54.43% | 4 |
| *N. linzhiensis* | 5 | 6 | 4 | 3 | 3 | 2 | 3 | 50%: 4;<br>50%: 3 | 3 | 102.88 | 32.25 | 31.35% | 8 |
| *N. bershulaensis* | 5 | 6 | 4 | 70%: 4;<br>30%: 3 | 3 | 3 | 3 | 4 | 3 | **108** | **53.5** | **49.50%** | **2** |

**Figure 3.**
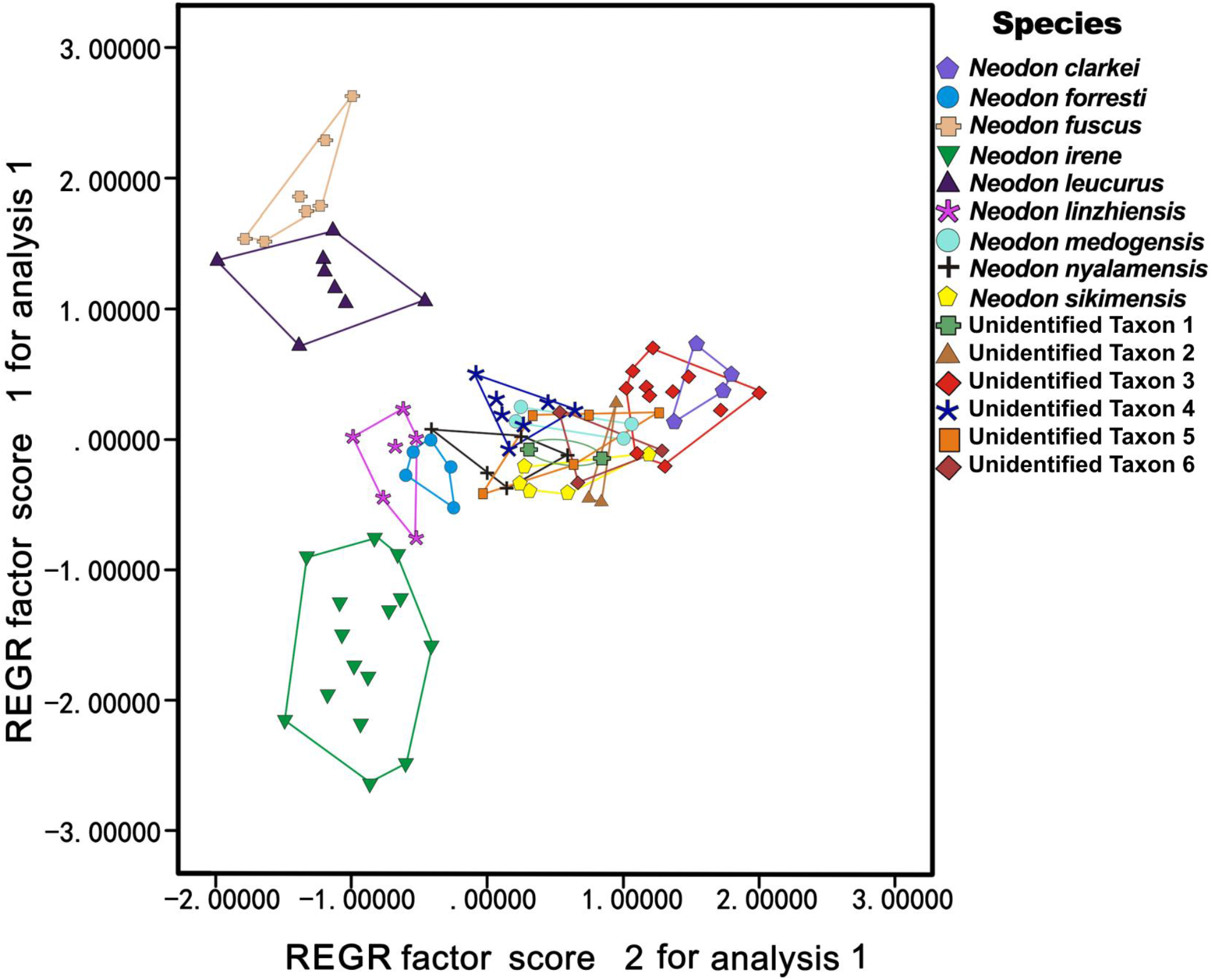
PCA result. Unidentified Taxon 1: unidentified taxon from Bershula Mountains; Unidentified Taxon 2: unidentified taxon from Chayu County; Unidentified Taxon 3: unidentified taxon from southern Namchabarwa Mountains; Unidentified Taxon 4: unidentified taxon from north of Yarlung Zangbo River; Unidentified Taxon 5: unidentified taxon distributed between south of Yarlung Zangbo River and north of Namchabarwa Mountains; Unidentified Taxon 6: unidentified taxon from Bomi County.

**Figure 4.**
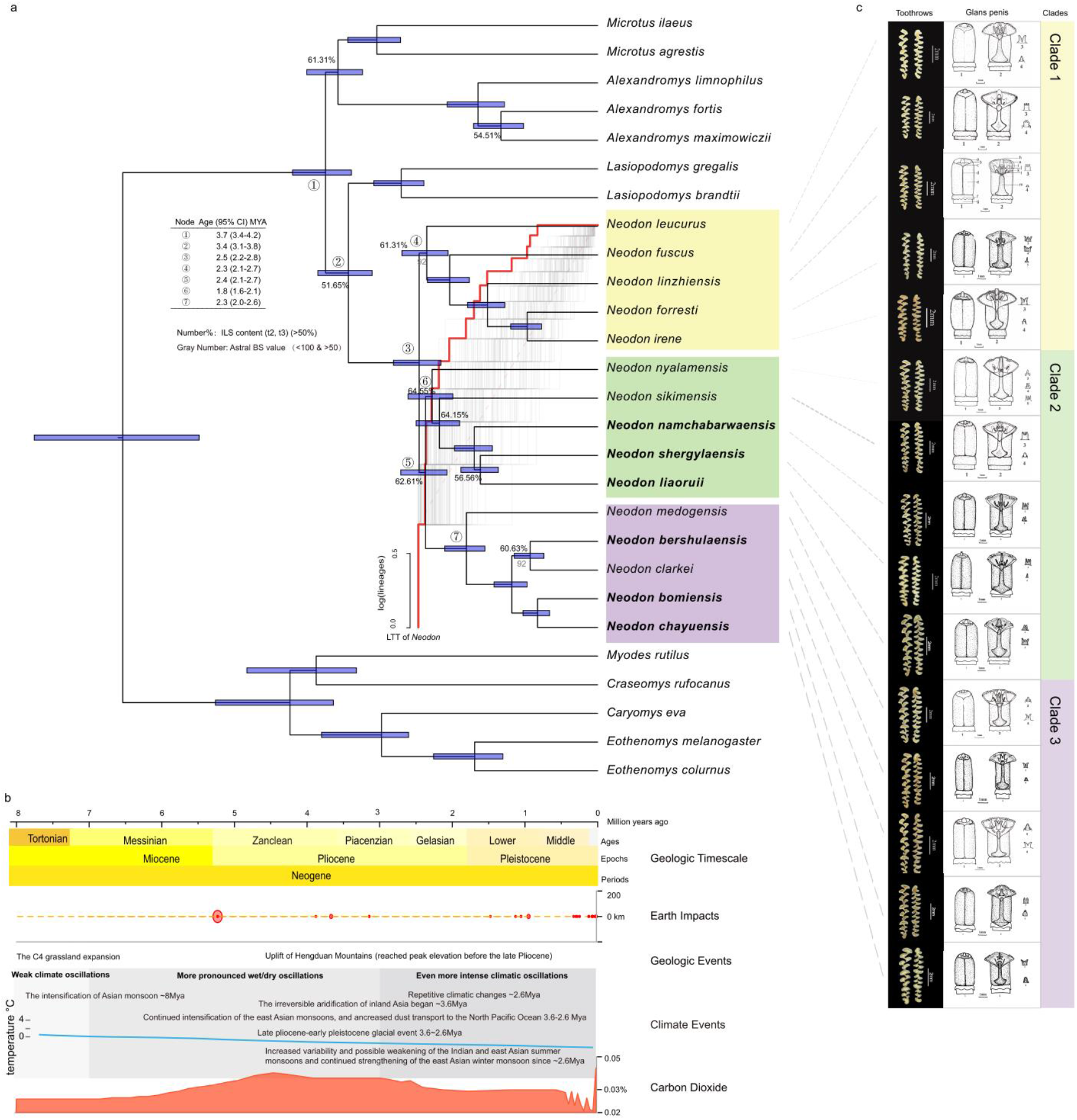
Divergence time tree, diversification patterns, ILS contents and morphologic photos for *Neodon*. a) Divergence time tree with the Astral branch supports number of nuclear gene tree were also showed near the branches. New species were marked in bold. Clades of *Neodon* were marked in shading with different colors on the tree (clade 1: yellow; clade 2: green; clade 3 purple). Branches with high ILS occurrence frequency (>50%) was marked with the ILS content number. Log-Lineage-through-time (LTT) plots for *Neodon* were estimated from the time-calibrated phylogeny of *Neodon* (red curve) and the simulated trees under a Yule model (gray curve), and the semilucent red dashed line indicates the null distribution under a Yule process. b) The divergence time, geologic timescale, earth impacts, carbon dioxide, geologic events and climate events were showed at the bottom of the figure. c) Comparison of the tooth rows and glans penis of the *Neodon* species were showed in the right of the figure. Numbered views are 1: glans; 2: midventral cut view; 3: urethral lappet; 4: dorsal papilla (detailed in Supplementary Fig. S2). For *N. linzhiensis*, lettered structural features are: a. distal baculum; b. outer crater; c. inner crater; d. ventral groove; e. glans; f. prepuce; g. penis body; h. station of dorsal papilla; i. lateral baculum (cartilage); j. urethral lappet; k. lateral baculum (bony part); l. distal baculum (bony part); and m. proximal baculum (Jia, et al. 2012).

#### PCA

Taxonomic identifications were corroborated by the PCA analysis (Fig. 3). The analysis included 17 non-gender-related measurements of external, cranial and dental characteristics of adults. Fitness testing delivered a Kaiser-Meyer-Olkin value of 0.941 and a Bartlett’s test of < 0.001; these demonstrated the robustness of the inference.

The first two principal components (PCs) explained 82.129% of the total variance. Thirteen measurements (LM, ZB, SGL, MB, SBL, SH, LIL, ABL, MM, CBL, LMxT, LMbT and HBL) contributed 60.653% to PC1, and four measurements (TL, IOW, EL and HFL) contributed 21.476% to PC2. Specimens of *N. irene*, *N. fuscus*, *N. leucurus* clearly differed from all other specimens. *Neodon forresti* overlapped slightly with *N. linzhiensis* and these two species clearly separated from remaining taxa. Specimens of *N. sikimensis* separated clearly from congeners from north of Yarlung Zangbo River and specimens from north of Yarlung Zangbo River separated unambiguously from specimens distributed south of Namchabarwa Mountains. Specimens from south of Yarlung Zangbo River differed from those north of Yarlung Zangbo River and specimens from south of this river differed with those from south of Namchabarwa Mountains. Specimens from south of Yarlung Zangbo River and *N. sikimensis* slightly overlapped with each other. Among specimens having four closed triangles in the first lower molar, *N. fuscus* and specimens from Chayu County clearly separated from each other and all other specimens. Specimens from Bomi County mixed with *N. medogensis*. For the specimens that have five closed triangles in the first lower molar, *N. clarkei*, *N. linzhiensis* and specimens from the Bershula Mountains separated from each other distinctively. ANOVAs for the scores of PC1 (F = 45.720, P < 0.001) and PC2 (F = 53.848, P < 0.001) exhibited highly significant differences among species of *Neodon*.

Tukey’s post hoc tests revealed that the PC1 or PC2 scores differentiated the six unidentified taxa. Among taxa that have 3 closed triangles on the first lower molar, the taxon from north of Yarlung Zangbo River differed significantly from the taxon from southern Namchabarwa Mountains (P < 0.001), *N. irene* (P < 0.001), *N. leucurus* (P < 0.001), and *N. forresti* (P=0.07), but did not differ significantly from the taxon from south of Yarlung Zangbo River, *N. sikimensis* and *N. nyalamensis*. The taxon from south of Yarlung Zangbo River also differed significantly from the taxon from Namchabarwa Mountains (P < 0.001), *N. irene* (P < 0.001), *N. leucurus* (P < 0.001), and *N. forresti* (P < 0.001), but not from the taxon from north of Yarlung Zangbo River, *N. sikimensis* and *N. nyalamensis*. The taxon from the southern Namchabarwa Mountains differed significantly of all taxa that have 3 closed triangles on the first lower molar. For the taxa that have 5 closed triangles in the first lower molar, the taxon from Bershula Mountains differed significantly from *N. linzhiensis* (P = 0.003) and *N. clarkei* (P < 0.001). For the taxa that have 4 closed triangles in the first lower molar, (a) the taxon from Chayu County differed significantly of *N. fuscus* (P < 0.001), but from the taxon from Bomi County and *N. medogensis*. The taxon from Bomi County also differed significantly from *N. fuscus* (P < 0.001). (4) For taxa that overlapped in PCA, *t*-tests were calculated and the results showed that at least 2 measurements had significant differences in one-to-one comparisons.

### HTS data

We obtained a total of 620.45 Gb clean paired-end reads for the 47 samples with each having a data size ranging from 2.21 Gb to 43.38 Gb (Supplementary Table S1). All samples achieved full mitochondrial genomes that contained 13 protein coding genes (PCGs) and two rRNAs for phylogenetic analyses. 6,016 full-length single-copy orthologous genes that spanned a total length of 288,415,615 nucleotides were obtained from *Peromyscus maniculatus* and used for nuclear gene dataset construction. After the removal of the low confidential genes, we obtained 5,382 coding genes (the “nuclear gene set”), with an average, maximal, minimal length of 1,879, 18,018 and 60 nt, respectively, for all the samples (Supplementary Fig. S3 and Table S4).

### Phylogenetic analysis

#### Sequence divergence and species delimitation

For the mitochondrial genes, the results from *Neodon cox1* and *cytb* showed an average of 0.94% similarity, 1.11% for intra-species genetic distance and 11.00%, 11.30% for inter-species distance, among which, *N. sikimensis* showed the greatest intra-species genetic distances with an average of 5.23% (*cox1*) and 5.20% (*cytb*). Individuals of the taxon from Bomi County and *N. forresti* did not show any genetic variation (Supplementary Table S5), and the smallest congenic inter-species genetic distances of 3.55% (*cox1*) and 3.70% (*cytb*) occurred between taxa from Bomi and Chayu counties (Supplementary Table S6-8, Supplementary Fig. S4). The results of molecular-based species delimitation agreed with the morphology analyses (Supplementary Appendix S1).

#### Phylogenetic Relationship

The phylogeny for species of *Neodon* was inferred using concatenated methods MrBayes and RAxML, and coalescent-based ASTRAL-III and SVDquartets (Fig4, Supplementary Fig. S5-8). Trees produced by all phylogenetic analyses based on nuclear genes yielded same topologies with several small-scale incongruences compared to that based on mitochondrial genes (Fig4, Supplementary Fig. S4-6). Both genomes resolved three clades. The species-tree inferred using ASTRAL III with a normalized quartet score of 70.13% and high branch support (BS ≥ 92) from the most comprehensive dataset of 5,328 nuclear genes was used for the following analyses; more details on the other trees were placed in Supplementary Fig. S5-8.

The first clade included five named taxa distributed in almost all of the HTR, especially in the eastern and western regions (Fig. 1 and 4, Supplementary Fig. S9a). The most wildly distributed species, *N. leucurus*, was the root taxon. The second clade included two described and three new taxa distributed mainly in the Himalayas (Supplementary Fig. S9b). *N. sikimensis* and *N. nyalamensis* were the root taxa and these specimens were collected near their type localities. The three new taxa occurred around the Yalung Zangbo River, Nachabarwa Mountains and Duoxiongla Peak area. Clade three, distributed in the eastern Himalayan and eastern HD mountains, included two described and three new species that were also collected from new sites: Bershula Mountains, Bomi County and Chayu County. Galongla Peak and the Gangrigabu Mountains separated the samples from Bomi and *N. medogensis* (Supplementary Fig. S9c).

### Species nomination

Comparisons of molar pattern and glans penes, principal component analysis of morphology, gene distance as well as the phylogenetic analyses all confirmed that six unidentified taxa were new species that were not described literally. We nominated them as *Neodon shergylaensis* sp. nov. (unidentified taxon from north of Yarlung Zangbo River), *Neodon namchabarwaensis* sp. nov. (unidentified taxon from between south of Yarlung Zangbo River and north of Namchabarwa Mountains), *Neodon liaoruii* sp. nov. (unidentified taxon from southern Namchabarwa Mountains), *Neodon chayuensis* sp. nov. (unidentified taxon from Chayu county), *Neodon bomiensis* sp. nov. (unidentified taxon from Bomi county), and *Neodon bershulaensis* sp. nov. (unidentified taxon from Bershula Mountains). Details of description could be found in Supplementary Appendix S1.

### Divergence time and diversification rate

Given the 95% credibility intervals of estimated date based on nuclear gene sets, the last common ancestor (LCA) of *Neodon* and outgroups lived in the late Miocene about 6.5 Mya (95% CI=7.7–5.5 Mya), and *Neodon* and *Lasiopodomys* diverged 3.4 Mya (3.8–3.1 Mya). *Neodon* experienced an explosive radiation in the late Neogene, and this coincided with changes in the Asian monsoons in the late Pliocene–early Pleistocene glacial event and uplifting events of the HD. The LTT plot suggested a deceleration in the rate of speciation after the initial radiation, and this was corroborated by the gamma statistic (γ= −3.6521, p < 0.05) (Fig. 4).

Although the three clades existed in different regions with different climates, the Kruskal-Wallis test of the substitution ratio did not show significant differences between them (Supplementary Fig. S10, Table S9-10) revealing that they still had similar evolutionary rates among genes and were not significantly influenced by the conditions of their distributions.

### Incomplete lineage sorting

Scans for the presence of ILS spanning the evolution of *Neodon* used 5,382 nuclear gene trees of 27 species (Supplementary Fig. S11a, b). ILS occurrence-frequency ranged from 1.49% to 64.55% for the 24 branches, while 37.50% of branches were affected by high levels of ILS (ILS rate > 50%) (Fig. 4). Thus, conflicting branches among gene trees were likely caused by high ILS content. A strong significant negative correlation was detected between the frequency of ILS occurrence and inner-node branch length (Pearson coefficient r = −0.98, P=6.12e-18) indicating that the high levels of ILS related to the rapid radiation (Supplementary Fig. S11c).

### Ancestral range reconstruction

The ancestral area estimation was conducted using BioGeoBEARS with the best model DEC+J detected by AIC testing. The results (Fig. 5), with a j value of 0.043 and d value of 0.012 (Supplementary Table S11), indicated that (1) the common ancestors of *Neodon* were located in the QTP, followed in some lineages by rapid radiations, coinciding with recent climate change and uplifts of the HD; (2) several clades exhibited dispersal events to other regions, and thus dispersed out of the QTP.

**Figure 5.**
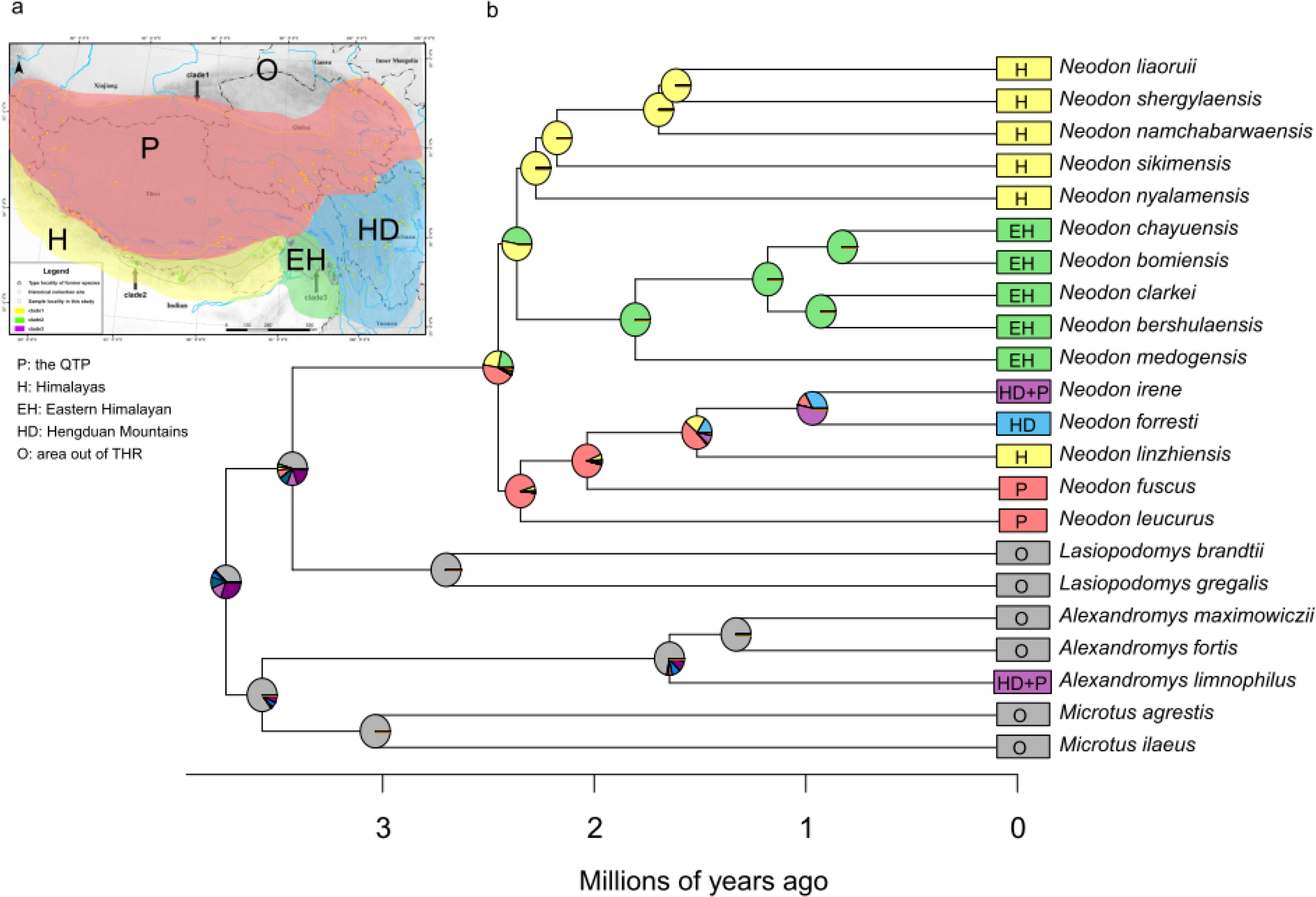
Ancestral range estimation results. a) Area delineation. b) Ancestral range estimation based on DEC + J model implemented in BioGeoBEARS.

## Discussion

Both the morphological and genetic data strongly support the long-term underestimated diversity in *Neodon*. Our sampling contributes six new species and confirms a total of 15 extant species of *Neodon*, which is much greater diversity than previously documented.

Resolution of the number of closed triangles in the first lower molar (CTFLM) was thought to identify species of *Neodon* (Feng, et al. 1986; Jia, et al. 2012; Luo 2000). However, our analyses show its failure to do so because, for example, all specimens with three CTFLM (distributed in southern Tibet) were once regarded as *N. sikimensis*, but now includes at least five species: *N. sikimensis, N. nyalamensis, N. linzhiensis, N. liaoruii* sp. nov., *N. shergylaensis* sp. nov., and *N. namchabarwaensis* sp. nov. The genetic distances between these species range from 7.61% to 13.17% for *cytb* and 8.01% to 12.74% for *cox1*. Analyses obtain the same result for four and five CTFLMs (detailed in Supplementary Table 5-8). In addition to CTFLM, our results show that several other traits are critical for classifying species of Arvicoloni (Jia, et al. 2012; Liu, et al. 2017) including external, cranial, dental and penis characters, which differ at various levels between species of *Neodon*, and especially glans penes and bacula (Fig. 4).

Our phylogenetic analyses take advantage of thousands of nuclear genes as well as mitochondrial protein-coding genes and it is by far the largest molecular dataset for the study for small mammals. The results using coalescent and concatenation methods show good consistency and corroborate *Lasiopodomys* as the sister group of *Neodon*. They share a common ancestor with a clade consisting of *Microtus* and *Alexandromys*. While species in *Neodon* form three clades, the newly discovered species occur mainly in clades two and three (Fig. 4). Speciation in *Neodon* appears to have occurred in a very short time and resulted in at least 15 species. The rapid speciation also likely led to the negative correlation between percentage of ILS and inner-node branch length of the dated tree.

Geographical isolation as a driver for speciation has been studied intensively in a myriad of taxa (Coyne and Orr 2004; Winger and Bates 2015; Xing and Ree 2017). Our results demonstrate that several critical geographical events during the orogenesis of HD and repetitive climate changes with glacial cycles play crucial roles in the diversification of these voles. Global cooling, irreversible aridification of inland Asia and uplifting episodes of the HD coincide with the divergence of *Neodon* and *Lasiopodomys*; subsequent climate change and dispersal events and of *Neodon* led to the adaptive radiation. The coincidence of *Neodon*’s evolutionary history with the several climate changes and orogenic events of HD reveal that the QTP is the origin and evolution center for *Neodon*. This corresponds with the “out of QTP” hypothesis. The two most widely distributed species of *Neodon*, *N. leucurus* and *N. fuscus*, occupy root positions within *Neodon*, confirming *Neodon*’s origin in the QTP. Further, climate oscillations and the formation of rivers and mountains on the HD gave rise to physical boundaries for dispersal of *Neodon*, and subsequent speciation. This fits well with the concept of “sky island effects” (Supplementary Fig. S9). For instance, at the eastern THR (including HD and the eastern margins of QTP and HM) where mountain ranges or single summits served as glacial period sky island refugia (Mosbrugger, et al. 2018), *N. namchabarwaensis* sp. nov. and *N. liaoruii* sp. nov. appear to have speciated due to the obstruction posed by the Duoxiongla Mountain pass (4,200 m above sea level (a.s.l.)), and *N. medogensis* and *N. bomiensis* differentiated due to the barrier effect of Galongla Snow Mountain pass (4,200 m a.s.l.) (Supplementary Appendix S1). Finally, systematic sampling and integrative approaches are needed for other relatively sedentary species, such as small mammals, reptiles, amphibians, and others in the THR, to clarify their biodiversity.

## SUPPLEMENTARY MATERIAL

Supplementary materials and data files are available from Dryad data repository doi. Trees conducted by this study also could be found in http://purl.org/phylo/treebase/phylows/study/TB2:S25507.

## ACKNOWLEDGMENTS

We thank Rui Liao for assistance in collecting specimens in the field. Special thanks to Yinjuan Mao and Junhua Bai for drawing the morphological figures.

