## Supplementary Appendix 1 for "Out of the Qinghai-Tibetan Plateau and get flourishing - the evolution of *Neodon* voles (Rodentia: Cricetidae) revealed by systematic sampling and low coverage whole genome sequencing"

SUPPORTING ONLINE MATERIAL FOR

### Out of the Qinghai-Tibetan Plateau and get flourishing - the evolution of *Neodon* voles (Rodentia: Cricetidae) revealed by systematic sampling and low coverage whole genome sequencing

SHAOYING LIU[1*#], CHENGRAN ZHOU[2, 3, 4#], TAO WAN[1, 5], GUANLIANG MENG[3, 4], W. ROBERT W. MURPHY[6], ZHENGXIN FAN[2], MINGKUN TANG[1], YANG LIU[1], TAO ZENG[2], SHUNDE CHEN[7], YUN ZHAO[2], SHANLIN LIU[3, 4, 8]*

#contribute equally to this paper

**THIS PDF FILE INCLUDES**

Materials and Methods

Results and Discussion

References

Keys for species Identification

Supplementary Figures Captions

Supplementary Tables Captions

List of Supplementary Appendices

#### **SUPPORTING ONLINE MATERIAL**

##### MATERIALS AND METHODS

*Gene data construction.*—We obtained orthologous genes for each sample via genome variance (SNP and InDel) identification and consensus calling function in bcftools v.1.8 with the following criteria: 1) minimum read number for indel candidates of 5; 2) properly paired reads; 3) optical duplicates and supplementary alignments skipped using default parameter; 4) SNP p-value of 1e-3 in –m mode; 5) filtering SNPs within 3 bp of an InDel and filtering clusters of InDels separated by 10 or fewer bp allowing only one to pass. In addition, we kept the reference allele for heterozygous sites and masked sites where the DP4 value (number of high quality mapped reads) was smaller than 1 to 'N'.

*Divergence time estimation.*—Divergence times on the species tree were estimated based on the nuclear genes using the MCMCTree implemented in the PAML v.4.9h package (Yang 2007) with the approximate likelihood calculation of 'REV' (GTR, model=7) model. First, second codon sites of the "high coverage nuclear gene set" were extracted to generate a new supergene and the final Astral tree was extracted as the input tree for subsequent dating analysis. Then, fossil calibration points were taken from the Paleobiology and timetree databases, setting a prior of divergence time for the nodes on the tree. The prior for mean substitution rate was estimated using BaseML. Gradient and Hessian matrices were obtained using MCMCTree with 'correlated rates clock' (clock=3), overall substitution rate (rgene_gamma) set at G (1, 15.7) and rate drift parameter (sigma2_gamma) at G (1,

4.5). Finally, MCMC analyses were run twice with a burn-in of 50,000 iterations and different random seed numbers to check the convergence. Each run was sampled every 5000 iterations until 10000 samples were gathered. Tracer (Rambaut, et al. 2018) was also used to examine the convergence of MCMC analysis results.

#### RESULTS AND DISCUSSION

##### *Sequence divergence and species delimitation*

*Sequence divergence*.—The ranges of minimum average nuclear genetic distances between the newly described taxa and existing species ranged from 0.17% (taxon from Bomi County and *N. clarkei*.) to 0.42% (taxon from southern Namchabarwa Mountains and *N. irene*, *N. sikimensis*, *N. medogensis* and *N. clarkei*). Within described species, *N. irene* and *N. forresti* had the minimum nuclear genetic distance of 0.18% (Supplementary Figure S4).

*Species delimitation*.—Results of molecular-based species delimitation agreed with the morphological analyses. The number of species in our sample was also estimated using ABGD with aligned mitochondrial gene set and bPTP with mitochondrial trees (Supplementary Fig. S6) and nuclear trees (Supplementary Fig. S7). The ABGD method obtained a stable result and delimited 29 species including outgroup taxa. The ABGD result supported most of the novel species groups in our samples, but one split occurred within *N. sikimensis*. For mitochondrial data set, the ML solution of bPTP delimited 29 species, which agreed with ABGD, while the BI solution of bPTP delimited 31 species, including two more splits in *N. forresti* and *N.*

*irene*. For nuclear data set, the ML solution of bPTP delimited all 16 putative species of *Neodon* while the BI solution of bPTP delimited 15 species of *Neodon*.

*Hypothesis of Speciation and Evolution of Neodon*

The time-calibrated genomic analysis (Fig. 5), combined with the current distribution (Fig. 1) and the geological and climatic events (Fig. 5), provided important clues for understanding evolutionary patterns in *Neodon*.

*The divergence of the common ancestor species of Neodon and Lasiopodomys.*—In the middle Pliocene, Earth’s surface generally cooled. Orogenesis in the THR created a large barrier effect on the ancient southwest monsoons and resulted in the occurrence of the South Branch Westerly Jetstream, directly, and further promoted the transformation of the environment on surface of the plateau into a humid environment (An, et al. 2001; Guo, et al. 2008). The common ancestor *Lasiopodomys* and *Neodon* began to differentiate as it adapted to different climates. One branch, which adapted to the drought and grassland environments, retreated to the arid north plateau and evolved into the *Lasiopodomys*, while another branch, which adapted to the humid environment, occupied the humid plateau surface and evolved into *Neodon*.

*The first divergence of Neodon.*—The oldest ancestors within *Neodon* involved *N. leucurus*, and they were distributed wildly on the plateau (Fig. 5). In the initial stage no barrier to the distribution of *N. leucurus* existed because the Yarlung Zangbo River has not yet formed (Wang, et al. 2002). Climatic changes and dispersal events caused the first divergence of the ancestral species and resulted into three evolutionary branches. The branch that includes *N. leucurus* and *N. fuscus* continued to occupy the plateau. Another branch existed on the southern edge of the QTP and evolved into *N.*

*nyalamensis* and *N. sikimensis*, which occur around the Himalayas. The third branch occupied the southeastern margin of the region and evolved into *N. medogensis*. The Yarlung Zangbo River formed and became a barrier to dispersal of *Neodon*, and then *N. leucurus* and *N. fuscus* occupied most of the northern part of the plateau while *N. nyalamensis* and *N. medogensis* occupied the narrow area on the south of the Yarlung Zangbo River separated by the Palung Zangbo Ancient River Channel in the eastern and western areas (Chen, et al. 2008).

*Rapid Adaptation of Neodon and the sky island species.*—At the early stage of rapid radiation, the Himalayas, Kangchenjunga Peak and Namcha Barwa Peak isolated the ancestors of the three major lineages. During the late Pliocene–early Pleistocene glacial stage, HD uplifted and the eastern margin of the plateau was formed. Glacial events caused the formation of large-scale ice sheets on the plateau's surface, and this drove some populations of *Neodon* to relatively warm and humid refugia in the southeastern plateau. These events and the barriers formed by mountains and the connection of the Yarlung Zangbo and Palong Zangbo rivers led to more speciation events, including the formation of *N. irene*, *N. forresti*, *N. clarkei*, *N. bershulaensis*, *N. chayuensis* and *N. bomiensis*.

*Neodon shergylaensis* and *N. linzhiensis* occur on both sides of Niyang River and Sejila Mountain pass (4,728 m a.s.l.) (Supplementary Fig. S1). This suggests that the multiple glacial obstruction of the Niyang River (Liu, et al. 2006) did not function as a barrier to dispersal for *Neodon*. However, the Yarlung Zangbo River (and ancient Palon Tsangpo River) separated *N. namchabarwaensis*, *N. shergylaensis* and *N. nyalamensis*. In addition, Duoxiongla Mountain (pass: 4,200 m a.s.l.) formed a barrier for *N. namchabarwaensis* sp. nov. and *N. liaoruii* sp. nov., and Galongla Snow Mountain (pass: 4,200 m a.s.l.) isolated *N. medogensis* and *N. bomiensis* (Supplementary Fig. S2). Thus, these constitute sky island species. Although *N. irene* evolved recently, it has a wide distribution in the eastern and southeastern parts of the plateau and it may have experienced a recent population expansion.

#### KEYS FOR SPECIES IDENTIFICATION

For identifying species of *Neodon*, we provide the following key:

1. First lower molar usually with 5 closed triangles (very few species with 4 closed triangles), distal and lateral bacula, at least the distal bacula long and sturdy: *Microtus*

   First lower molar with 5, 4 or 3 closed triangles; the distal baculum very short, the lateral bacula only slightly ossified or cartilage: *Neodon* .............................................. 2

2. First lower molar with 3 closed triangles .................................................. 3

   First lower molar with 4 or 5 closed triangles .................................................. 10

3. TL/HBL larger than 50% in average .............................................. *N. liaoruii* sp. nov.

   TL/HBL less than 50% in average .................................................. 4

4. HBL less than 100mm .................................................. *N. irene*

   HBL larger than 100mm .................................................. 5

5. Most of specimens with 4 inner and 4 outer angles in the third upper molar ........... .................................................. *N. nyalamensis*

   The third upper molar with 4 or 3 inner and 3 outer angles .................................. 6

6. The first lower molar with 5 inner and 3 outer angles ................................ *N. leucurus*

   Labial side of the first lower molar with 4 or 5 angles .............................................. 7

7. The second upper molar with 2 inner and 3 outer angles ............................ *N. forresti*

   The second upper molar with 3 inner and 3 outer angles .......................................... 8

8. The first upper molar with 4 inner and 3 outer angles .................................................. .................................................. *N. namchabarwaensis* sp. nov.

The first upper molar with 3 inner and 3 outer angles............................................9

9. TL/HBL approximately 44%..................................................................*N. sikimensis*

TL/HBL less than 38% in average......................................*N. shergylaensis* sp. nov.

10. The first lower molar with 4 closed triangles .......................................................11

The first lower molar with 5 closed triangles .......................................................14

11. The second upper molar with 2 inner and 3 outer angles ........................... *N. fuscus*

The second upper molar with 3 inner and 3 outer angles ......................................12

12. The third upper molar usually with 4 inner and 3 outer angles ..........*N. medogensis*

The third upper molar with 3 inner and 3 outer angles..........................................13

13. The first upper molar of majority of specimens with 4 inner and 3 outer angles...... ........................................................................................*N. chayuensis* sp. nov.

The first upper molar with 3 inner and 3 outer angles...............*N. bomiensis* sp. nov.

14. TL/HBL larger than 50%............................................................................*N. clarkei*

TL/HBL less than 50% ..........................................................................................15

15. The second upper molar with 2 inner and 3 outer angles .................... *N. linzhiensis*

The second upper molar with 3 inner and 3 outer angles......................................... ..........................................................................*N. bershulaensis* sp. nov.

**Family Cricetidae Rochebrune, 1883**

**Subfamily Arvicolinae Miller, 1906**

**Genus *Neodon* Horsfield, 1841**

***Neodon shergylaensis* sp. nov.**

**Shergyla Mountain vole**

*Holotype*.—Adult male, field number XZGB09N195 (Museum number: SAF091027), collected by Liao Rui and Liu Yang on 30 May 2009. Specimen preserved at the Sichuan Academy of Forestry as a skin, cleaned skull, penis and tissues. External and cranial measurements (in mm) as follows (abbreviations in Material and Methods): HBL117.0 mm; TL 41.0 mm; HFL 20.0 mm; EL 15.0 mm; SGL 28.22 mm; SBL 26.41 mm; CBL 27.41 mm; ZB 15.79 mm; IOW 4.13 mm; MB 12.56 mm; SH 10.60 mm; ABL7.96 mm; LMxT 6.32 mm; LMbT 6.01 mm; LM 19.55; M-M 5.67 mm; and OLLI 8.66 mm. Body mass 44 g. The photos of skull, dentition, and mandible are in Supplementary Fig. S1a.

*Type locality*.—Shergyla Mountains, southeast of Xizang, China, 29.62368° E, 94.66174° N, elevation 4500 m. This specimen was caught with a steel trap (Jiangxi Mouse Devices Factory (JMDF)) in shrubs of *Salix cupularis*, *Rhododendron alutaceum* and *Sabina wallichiana*.

*Paratypes*.—Thirty specimens (14 males and 16 females). Nine intact topotype adults (3♂♂, 6♀♀), field numbers: LZRAP01013♂, LZRAP01017♂, LZRAP01020♀, LZRAP01014♀, LZRAP01019♀, XZGB09N197♀, XZGB09N 262♂, LZRAP01024♀, GB0815001J♀); five topotype adults with skulls broken (2♂♂, 3♀♀), field numbers: XZPAR01016♀, XZPAR01023♂, XZBG0802001♂, XZGB09N185♂, XZGB09N186♀; 16 juveniles (6 intact, 10 with skulls broken; 8♂♂, 8♀♀), field numbers: LZRAP01010♂, LZRAP01011♀, LZRAP01015♀, XZGB09N196♀, DJ01001♀, XZGB0802002♀, LZRAP01018♂, LZRAP01021♂,

LZRAP01022♀, LZRAP01025♂, XZGB0801001♀, XZGB0801002♂, XZGB0802001♂, XZGB0816001♂, XZGB09N187♀, XZGB09N198♂.

*Distribution*.—Known from north of the Yarlung Zangbo River at over 3160m a.s.l. both sides of Shergyla Mountains and Niyang River.

*Etymology*.—The species is named for its type locality. This region supports a high biodiversity. In addition to *N. shergylaensis* sp. nov., *N. linzhiensis* occurs in lower regions of the same mountain. *Ochotona macrotis* occurs in the same habitat of this new species. The species' epithet highlights the importance of conserving this area's endemic biodiversity.

*Diagnosis*.—Medium body, average 115.7 mm in length (adult); hind feet 19.5 mm in average. Tail length 42.7 mm in averages, approximately 37% of HBL. The first lower molar with 3 closed triangles, 6 inner and 4 outer angles in 64% specimens, 6 inner and 5 outer angles in 36% specimens. 1$^{st}$ upper molar with 3 inner and 3 outer angles. 2$^{nd}$ upper molar 3 inner and 3 outer angles. 3$^{rd}$ upper molar with 4 inner and 3 outer angles (Supplementary Fig. S1a).

*Description*.—Pelage from head to hip uniform brown-black. Entire back covered with fine, dense, velvet hair. Ventral hairs with gray-black base and gray-white tip. Transition between dorsal and ventral pelage vague. Ears project slightly above pelage, covered with dense gray-black hairs. Tail bicolor obvious, dorsal tail brown-black, ventral tail grey-white; hairs on top of tail slightly longer. Dorsal surface forefoot and hindfoot yellow-white. Claws yellow-white. Five palmar and 5 plantar pads. Females with 1 pair inguinal and 1 pair pectoral mammae.

Skull sturdy, in dorsal profile straight and brain case flattened. Nasal broad anteriorly narrowing posteriorly. Parietal elliptic and a protrusion on the side.

Interparietal broad, anterior part triangle-shaped and posterior margin arc-shaped (Supplementary Fig. S1a). Interorbital and temporal ridges absent. Zygomatic arches slender and middle part slightly broader. Auditory bullae moderately sized. Incisory foramen short and narrow. Posterior palate typical of *Microtus*, with 2 obvious lateral pits. Many small foramens in palatine. Mandibles medium-sized (Supplementary Fig. S1a).

Upper incisors orange. Molars rootless. $1^{st}$ upper molar with 4 closed triangles after the anterior transverse space, 3 outer and 3 inner angles. $2^{nd}$ upper molar with 3 closed tooth rings after the anterior transverse space, forming 3 inner and 3 outer angles. $3^{rd}$ upper molar without closed tooth rings with 4 inner and 3 outer angles (Supplementary Fig. S1a6). $1^{st}$ lower molar with 3 closed triangles, holotype with 6 inner and 4 outer angles, but 36% of type series with 6 inner and 5 outer angles. $2^{nd}$ and $3^{rd}$ lower molars with 3 outer and 3 inner angles (Supplementary Fig. S1a7).

Glans penis (Supplementary Fig. S2h) pole-like and slender with a ventral groove. Outer crater no obvious papilla. Urethral lappet with 3 forks. Dorsal papilla with 1 tip. Proximal baculum bony with a rhombus-shaped base. Distal baculum also bony and tongue-shaped. Lateral bacular processes bony, stick-shaped and very short.

*Reproduction*.—In late May and early of June, most adult males orchidoptosis, but no females pregnant. In mid-September, over 50% adult females pregnant, with 1–3 embryos, but no male orchidoptosis. No data exist for other months.

*Habitat*.—This species inhabits fir forest at elevations higher than 3800 m a.s.l., fir height approximately 18 m, 70% coverage. Understory, humus 10–20 cm thick. Moss very abundant, 80% coverage and many fallen dead trees.

**Family Cricetidae Rochebrune, 1883**

#### Subfamily Arvicolinae Miller, 1906

#### Genus *Neodon* Horsfield, 1841

#### *Neodon namchabarwaensis* sp. nov.

#### Namchabarwa Mountain vole

*Holotype*.—Adult male, field number XZGB0818009 (Museum number: SAF08939), collected by Liao Rui on 30 May 2009. Specimen preserved at the Sichuan Academy of Forestry as a skin, cleaned skull, penis and tissues. The external and cranial measurements (in mm) as follows (abbreviations in Material and Methods): HBL121.0 mm; TL51.0 mm; HFL 21.0 mm; EL 15.0 mm; SGL 27.88 mm; SBL 26.74 mm; CBL 27.70 mm; ZB 15.36 mm; IOW 3.62 mm; MB 12.46 mm; SH 10.60 mm; ABL7.36mm; LMxT 6.12 mm; LMbT 6.44mm; LM 19.87; M-M 5.48 mm; and OLLI 9.18 mm. Body mass 44g. Photos of skull, dentition and mandible in Supplementary Fig. S1b.

*Type locality*.—Nanyi township, Milin County, south of Xizang, China, 29.17889° E, 94.15113° N, elevation 3160 m a.s.l. This specimen was trapped with a steel trap (Jiangxi Mouse Devices Factory) in secondary fir forest. Its habitat is a random riprap with thick humus.

*Paratypes*.—Twenty eight specimens (9 males and 19 females). Sis adult intact topotypes (3♂♂, 3♀♀), field numbers: XZGB0817007♂, XZGB0818006♀, XZGB0818008♀, XZGB08010♂, XZGB0828001♀, XZGB09N235♂; ten topotype adults with skulls broken (3♂♂, 7♀♀), field numbers: XZGB0817006♂, XZGB0818009♀, XZGB0819001♂, XZGB0820001♂, XZGB0820002♀, XZGB0820003♀, XZGB09N212♀, XZGB09N213♀, XZGB09N214♀, XZGB09N234; 12 juveniles (6 intact, 6 with skulls broken; 3♂♂, 9♀♀), field numbers: XZGB0817008♀, XZGB0818007♀, XZGB0818011♂, XZGB0819002♀, XZGB0819003♀, XZGB0820004♂, XZGB0820005♀, XZGB0820006♀, XZGB0821001♀, XZGB0821002♀, XZGB0821003♂, XZGB0821004♀.

*Distribution*.—Known from south of the Yarlung Zangbo River, north of the Himalayan Mountains and east of Shigatse. The lowest elevation is 3160 m a.s.l.

*Etymology*.—Species is named for famous Namcha Barwa Mountain, the highest mountain of this region where the new species occurs. This region has a high biodiversity, but serious human disturbance, which puts high pressure on its biodiversity. In addition to *N. namchabarwaensis* sp. nov., the recently described pika (*Ochotona yarlungensis*) also occurs there. The species epithet highlights the importance conserving this area's endemic biodiversity.

*Diagnosis*.—Medium body, average adult length 114.9 mm; hind feet average 20.1 mm. Tail length averages 46.4mm, approximately 40.4% of HBL. First lower molar with 3 closed triangles, 6 inner and 5 outer angles. 1$^{st}$ upper molar with 4 inner and 3 outer angles. 2$^{nd}$ upper molar 3 inner and 3 outer angles. 3$^{rd}$ upper molar with 4 inner and 3-4 outer angles (Supplementary Fig. S1b).

*Description*.—Appearance same as *N. shergylaensis*. Pelage from head to hip uniform brown-black. Entire back covered with fine, dense, velvet hair. Ventral black-grey, hairs with black base and a very small proportion of yellow-white tip. Transition between dorsal and ventral pelage vague. Ears project above pelage, covered with short gray-black hairs. Color dorsal tail black and ventral tail lighter; hairs on top of tail slightly longer. Dorsal surface of forefoot and hindfoot yellow-brown. Claws yellow-white. Five palmar and 5 plantar pads. Females with 1 pair inguinal and pectoral mammae.

Skull sturdy, in dorsal profile straight and brain case flattened (Supplementary Fig. S1b). Nasal broad anteriorly narrowing posteriorly. Parietal elliptic protruding laterally. Interparietal broad, rectangular, middle of anterior part protruding forward. Interorbital and temporal ridges presented. Zygomatic arches slender and middle part slightly broader. Auditory bullae moderately sized. Incisory foramen short and narrow. Posterior palate typical of *Microtus*, with 2 obvious lateral pits. Foramen in palatine very rare. Mandibles medium-sized.

Upper incisors orange. Molars rootless. 1$^{st}$ upper molar with 4 closed triangles after the anterior transverse space, and the last one protruding lingual forward, 3 outer and 4 inner angles. 2$^{nd}$ upper molar with 3 tooth rings and a posterior-interior small tooth ring after the anterior transverse space, forming 3 inner and 3 outer angles. 3$^{rd}$ upper molar with 3 closed triangles and a “C” tooth loop after the anterior transverse space, this tooth with 4 inner and 3-4 outer angles (Supplementary Fig. S1b6). 1$^{st}$ lower molar with 3 closed triangles and a semicircular anterior tooth cap; this tooth with 6 inner and 5 outer angles. 2$^{nd}$ and 3$^{rd}$ lower molars with 3 outer and 3 inner angles (Supplementary Fig. S1b7).

Glans penis (Fig. 4, Supplementary Fig. S2b) pole-like and slender with ventral groove. Outer crater with 4 obvious papilla on both sides. Urethral lappet with 3 forks. Dorsal papilla with 2 tip. Proximal baculum bony with a rhombus-shaped base, and the anterior bulged. Distal baculum also bony and dagger-shaped. Lateral bacular processes bony, bending and relatively longer.

*Reproduction*.—In early June, most adult males orchidoptosis, but females not pregnant. In mid-August, approximately 25% adult females pregnant, with 4 embryos, and 50% males orchidoptosis. No other data on reproduction are available.

*Habitat*.—This species inhabits spruce and fir forest at elevations 3160–3700 m a.s.l., tree height approximately 18 m, 50% coverage. Shrubs 2–3m and with 20% coverage. Understory, humus 5–10 cm thick and grass 10 cm, 40% coverage.

**Family Cricetidae Rochebrune, 1883**

**Subfamily Arvicolinae Miller, 1906**

**Genus *Neodon* Horsfield, 1841**

***Neodon liaoruii* sp. nov.**

**Liao's Mountain vole**

*Holotype.*—Adult male, field number XZ11117 (Museum number: SAF11302), collected by Liao Rui on 1 November 2011. Specimen preserved at the Sichuan Academy of Forestry as a skin, cleaned skull, penis and tissues. External and cranial measurements (in mm) as follows (abbreviations in Material and Methods): HBL120.0 mm; TL56.0 mm; HFL 21.0 mm; EL 14.0 mm; SGL 29.71 mm; SBL 27.02 mm; CBL 28.67 mm; ZB 15.7 mm; IOW 4.36 mm; MB 12.97 mm; SH 10.44 mm; ABL7.85 mm; LMxT 6.57 mm; LMbT 6.6 mm; LM 20.29; M-M 5.52 mm; and OLLI 9.09 mm. Body mass 42.74 g. Photos of skull, dentition and mandible in Supplementary Fig. S1c.

*Type locality.*—Nanyi township of Milin County, south of Xizang, China, 29.47028° E, 94.984° N, elevation 3260 m. Specimen caught with a steel trap (Jiangxi Mouse Devices Factory) in mixed coniferous broad leaved forest. Trap site with fallen deadwood and thick humus.

*Paratypes*.—50 specimens (24 males, and 26females). Ten intact adults (4♂♂, 6♀♀), field numbers: MT11036♀, MT11066♀, MT11067♀, MT11109♂, MT11118♂, MT11120♂, MT11122♀, MT11142♂, MT11143♀, MT11144♀.19 adults with skulls broken (9♂♂, 10♀♀), fields numbers: MT11032♂, MT11033♂, MT11034♀, MT11035♀, MT11037♀, MT11054♀, MT11062♂, MT11063♂, MT11065♂, MT11068♀, MT11082♀, MT11083♀, MT11084♀, MT11092♂, MT11094♂, MT11099♀, MT11107♂, MT11145♂, MT11146♀. Twenty-one juveniles (10 ♂♂, 11♀♀; 7 intact, 14 with skulls broken), field numbers: MT11064♂, MT11069♂, MT11070♂, MT11085♀, MT11093♂, MT11095♂, MT11096♂, MT11097♂, MT11098♀, MT11100♀, MT11101♀, MT11108♂, MT11110♀, MT11111♀, MT11112♀, MT11119♂, MT11121♂, MT11123♀, MM11124♀, MT11125♀, MT11126♀.

*Distribution*.—Known from south of the Yarlung Zangbo River, north of the Himalayan Mountains and east of Shigatse. Lowest elevation 2660 m a.s.l.

*Etymology*.—Species epithet is a patronym for the collector, Mr. Liao Rui. He made an important contribution on our collecting specimens. In the past 13 years, he has been to 25 provinces of China and collected over 10,000 specimens of small mammals, including ten new species.

*Diagnosis*.—Relatively large body, average 116.8 mm in length (adult); hind feet average 21.1 mm. Tail length averages 59.3mm, approximately 50.8% of HBL. First lower molar with 3 closed triangles, 6 inner and 5 outer angles. 1$^{st}$ upper molar with 3 inner and 3 outer angles. 2$^{nd}$ upper molar with 2 inner and 3 outer angles in 67% specimens, and 3 inner and 3 outer angles in another 33% specimens. 3$^{rd}$ upper molar with 4 inner and 3 outer angles in 61% specimens, and 3 inner and 3 outer angles in another 39% specimens (Supplementary Fig. S1c).

*Description*.—Appearance close to *N. namchabarwaensis*. Pelage from head to hip uniform brown-black. Entire back covered with fine, dense, velvet hair. Ventral hairs yellow-brown with black base. Transition between dorsal and ventral pelage vague. Ears project above pelage, covered with short yellow-brown hairs. Color of back of tail black and ventral of tail lighter; hairs at the top of tail slightly longer. Dorsal surface of forefoot and hindfoot grey-black. Claws yellow-white. Five palmar and 6 plantar pads. Females with 1 pair of inguinal and pectoral mammae.

Skull sturdy, in dorsal profile straight and brain case bulging slightly (Supplementary Fig. S1c1). Nasal broad anteriorly narrowing posteriorly. Parietal trapeziform with a lateral protrusion. Interparietal broad, irregular pentagon (Supplementary Fig. S1c1). Interorbital and temporal ridges presented but weaker. Zygomatic arches slender and middle part slightly broader. Auditory bullae moderately sized. Incisory foramen relatively long and broad. Posterior palate typical of *Microtus*, with 2 obvious lateral pits. Pterygoid with some foramina(Supplementary Fig. S1c2). Mandibles sturdy(Supplementary Fig. S1c5).

Upper incisors orange. Molars rootless. 1$^{st}$ upper molar with 4 closed triangles after the anterior transverse space, 3 outer and 3 inner angles. 2$^{nd}$ upper molar of holotype with 3 closed triangles forming 3 inner and 3 outer angles, but 66% specimens with 2 inner and 3 outer angles. 3$^{rd}$ upper molar of holotype with 4 inner and 3 outer angles, fourth outer angle vestigial; 39% specimens with 3 inner and 3 outer angles. 1$^{st}$ lower molar (Supplementary Fig. S1c6) with 3 closed triangles and a semicircular anterior tooth cap, tooth with 6 inner and 5 outer angles. 2$^{nd}$ and 3$^{rd}$ lower molars with 3 outer and 3 inner angles (Supplementary Fig. S1c7).

Glans penis (Fig. 4, Supplementary Fig. S2c) sturdy, pole-like and with a ventral groove. Outer crater with 3 obvious papilla on both sides. Urethral lappet with 3 forks, middle one slightly shorter. Dorsal papilla with 2 tips. Proximal baculum bony with a rhombus-shaped base, and concavely forward. Distal baculum bony, tongue-shaped. Lateral bacular processes bony and very short.

*Reproduction*.—In late October, approximately 10% adult males orchidoptosis, but females not pregnant. No other data on reproduction exist.

*Habitat*.—Species inhabits mixed coniferous broad leaved forest at elevations about 2700 m a.s.l., tree height approximately 18 m, 40% coverage. Shrubs 4 m and with 20% coverage. Understory, humus 5–10 cm thick and grass 50 cm, 50% coverage.

**Family Cricetidae Rochebrune, 1883**

**Subfamily Arvicolinae Miller, 1906**

**Genus *Neodon* Horsfield, 1841**

***Neodon chayuensis* sp. nov.**

**Chayu Mountain vole**

*Holotype*.—Adult female, field number CY37 (Museum number: SAF007607), collected by Liu Yang on 8 October 2007. Specimen preserved at the Sichuan Academy of Forestry as a skin, cleaned skull, and tissues. External and cranial measurements (in mm) as follows (abbreviations in Material and Methods): HBL109.0 mm; TL47.0 mm; HFL 20.0 mm; EL 16.5 mm; SGL 28.01 mm; SBL 26.92 mm; CBL 27.77 mm; ZB 16.46 mm; IOW 3.75 mm; MB 13.02 mm; SH 10.85 mm; ABL7.99 mm; LMxT 6.34 mm; LMbT 6.50 mm; LM 19.95; M-M 5.75 mm; and

OLLI 8.97 mm. Body mass 44g. Photos of skull, dentition, and mandible in (Supplementary Fig. S1d).

*Type locality*.—Chibagou National Nature Reserve, Chayu County, southeast of Xizang, China, 96. 98858° E, 28. 85716° N, elevation 2960 m a.s.l. Specimen was caught with a steel trap (Jiangxi Mouse Devices Factory) in a small valley, marsh wetland with dense of grass and thick moss.

*Paratypes*.—10 specimens (4 males, and 6 females). Three intact adults (1♂, 2♀♀), field numbers: CY35, CY44, CY45; 5 adults with skulls broken (2♂♂, 3♀♀), field numbers: CY36♀, CY38♂, CY47♂, CY48♀, CY49♀; 2 juveniles, field numbers: CY46♀, CY50♂.

*Distribution*.—Known from Cibagou National Nature Reserve, Chayu County, southeast of Xizang.

*Etymology*.—Species epithet derived from the county where collected. The biodiversity of Chayu County is under very high pressure. The name highlights the importance of conserving the endemic biodiversity of this area.

*Diagnosis*.—Medium body, average length 106.6 mm (adult); hind feet 19–21 mm. Average tail length 47. mm, approximately 45% of HBL. Tooth row sturdy. First lower molar with 4 closed triangles, but more or less confluent each other in many specimens, this tooth with 6 inner and 5 outer angles in 55% specimens; other 45% specimens with 6 inner and 4 outer angles. 1$^{st}$ upper molar with 4 inner and 3 outer angles in 67% specimens, another 33% with 3 inner and 3 outer angles. 2$^{nd}$ upper molar with 3 inner and 3 outer angles. 3$^{rd}$ upper molar with 4 inner and 3 outer angles (Supplementary Fig. S1d).

*Description*.—Pelage from head to hip uniform grey-brown. Entire back covered with fine, dense, velvet hair. Ventral hairs yellow-white with black base. Transition between dorsal and ventral pelage vague. Ears project above pelage slightly, covered with short grey-brown hairs. Tail obviously bicolored. Dorsal color of tail grey-black, ventral yellow-white; hairs at the top of tail slightly longer. Dorsal surface of forefoot and hindfoot grey-black. Claws grey-white, but upper surface of nail black-grey. Five palmar and 5 plantar pads. Females with 1 pair of inguinal and pectoral mammae.

Skull sturdy, in dorsal profile straight and flatten (Supplementary Fig. S1d). Nasal relatively short, broad anteriorly narrowing posteriorly. Parietal irregularly shaped with a lateral protrusion. Interparietal broad, irregularly shaped and the anterior part protruding forward. Interorbital and temporal ridges presented and relatively well-developed. Zygomatic arches relatively sturdy and middle part slightly broader (Supplementary Fig. S1d3). Auditory bullae moderately sized. Incisory foramen relatively long and broad. Posterior palate typical of *Microtus*, with 2 obvious lateral pits. Many foramina in palate pterygoid. Mandibles sturdy (Supplementary Fig. S1d5).

Upper incisors orange, sturdy. Molars rootless. Tooth row sturdy. 1$^{st}$ upper molar with 4 closed triangles after the anterior transverse space, 3 outer and 4 inner angles in holotype; 33% specimens with 3 outer and 3 inner angles. 2$^{nd}$ upper molar with 3 closed triangles forming 3 inner and 3 outer angles. 3$^{rd}$ upper molar without closed triangles, with 4 inner and 3 outer angles (Supplementary Fig. S1d2, 6). 1$^{st}$ lower molar of holotype with 4 closed triangles and a trilobal anterior tooth cap, which has 6 inner and 5 outer angles; other 45% specimens with 6 inner and 4 outer angles. In many specimens, triangles of the 1$^{st}$ lower molar are not closed entirely, more or less confluent each other. 2nd and 3rd lower molars with 3 outer and 3 inner angles (Supplementary Fig. S1d7).

Glans penis (Supplementary Fig. S2d) relatively sturdy and short, pole-like and with a ventral groove. Outer crater with 5 obvious papilla on both sides. Urethral lappet with 3 forks, middle one very short. Dorsal papilla with single tip and sturdy. Proximal baculum bony with rhombic base; middle of bottom concave upward. Distal baculum stick-shaped and pointed. Lateral bacular processes cartilaginous and long.

*Reproduction*.—In early October, approximately 20% adult males orchidoptosis; females not pregnant. No data are available for other months.

*Habitat*.—This species inhabits marshland at elevations about 3000 m a.s.l. Shrubs 2–3m and with 10% coverage. Grass 40—60 cm, 95% coverage. Humus 10–20 cm thick.

#### Family Cricetidae Rochebrune, 1883

#### Subfamily Arvicolinae Miller, 1906

#### Genus *Neodon* Horsfield, 1841

#### *Neodon bomiensis* sp. nov.

#### Bomi Mountain vole

*Holotype*.—Adult male, field number XZ13015 (Museum number: SAF13477), collected by Liao Rui on 31 October 2013. Specimen preserved at the Sichuan Academy of Forestry as a skin, cleaned skull, penis and tissues. External and cranial measurements (in mm) as follows (abbreviations in Material and Methods): HBL116.0 mm; TL53.0 mm; HFL 18.0 mm; EL 13.0 mm; SGL 27.56 mm; SBL 25.87 mm; CBL 26.81 mm; ZB 15.89 mm; IOW 4.35 mm; MB 12.53 mm; SH 9.90

mm; ABL7.51 mm; LMxT 6.48 mm; LMbT 6.52 mm; LM 19.96; M-M 5.625 mm; and OLLI 9.10 mm. Body mass 40g. Photos of skull, dentition, and mandible in (Supplementary Fig. S1e).

*Type locality*.—Bomi County, southeast of Xizang, China, 95.9575816° E, 29.82959° N, elevation 2900 m a.s.l. Specimen captured with a steel trap (Jiangxi Mouse Devices Factory) under a tall broad leaf tree set in dense grass and thick humus.

*Paratypes*.—4 specimens (4 females); 2 intact adults, field numbers: MT11304♀, MT11305♀; two specimens with skulls broken, field numbers: XZ13016♀, adult; XZ13031♀, juvenile.

*Distribution*.—Known only from the type locality, Bomi County, southeast of Xizang.

*Etymology*.—Species epithet derived from the county where type series collected. The biodiversity of Bomi County is under very high pressure. The name highlights the importance of conserving this area's endemic biodiversity.

*Diagnosis*.—Medium body, average length 111.75 mm (adult); hind feet 18–19mm (average 18.75). Tail length 53–56 mm (average 53.75 mm), approximately 48.1% of HBL. First lower molar with 4 closed triangles, 6 inner and 4 outer angles in 60% specimens; other 40% with 5 inner and 4 outer angles. 1st upper molar with 4 closed triangles, forming 3 inner and 3 outer angles. 2nd upper molar with 3 inner and 3 outer angles. 3rd upper molar with 4 inner and 3 outer angles (Supplementary Fig. S1e)

*Description*.—Pelage from head to hip uniform grey-black, brushed with brown-grey tinge. Entire back covered with fine, dense, velvet hair. Ventral hairs grey-white with black base. Transition between dorsal and ventral pelage vague. Ears project above pelage slightly, covered with short grey-black hairs. Tail single color. Dorsal tail black and ventral tail slightly lighter; dorsal tail hairs slightly longer. Dorsal surface of forefoot and hindfoot grey-black. Claws yellow-brown; upper surface of nails grey-black. Five palmar and 5 plantar pads. Females with 1 pair of inguinal and pectoral mammae.

Dorsal profile of skull straight; brain case bulging slightly (Supplementary Fig. S1e). Nasal relatively short, broad anteriorly narrowing posteriorly. Parietal irregular with lateral protrusion. Interparietal broad, irregular elliptic, mid-anterior part protruding forward. Interorbital ridges absent, temporal ridges present but weak. Zygomatic arches slender and middle part slightly broader. Auditory bullae moderately sized. Incisory foramen relatively longer and broader. Posterior palate typical of *Microtus*, with 2 obvious lateral pits. Many mini-foramen in palate and pterygoid. Mandibles medium (Supplementary Fig. S1e).

Upper incisors orange. Molars rootless. $1^{st}$ upper molar with 4 closed triangles after the anterior transverse space, 3 outer and 3 inner angles. $2^{nd}$ upper molar of holotype with 3 closed triangles forming 3 inner and 3 outer angles. $3^{rd}$ upper molar with 4 inner and 3 outer angles (Supplementary Fig. S1e6). $1^{st}$ lower molar of holotype with 4 closed triangles and a trefoil anterior tooth cap; this tooth with 6 inner and 4 outer angles, but 40% specimens with 5 inner and 4 outer angles. $2^{nd}$ and $3^{rd}$ lower molars with 3 outer and 3 inner angles (Supplementary Fig. S1e7).

Glans penis (Supplementary Fig. S2e) medium, pole-like, with a ventral groove. Outer crater with 10 obvious papilla on both sides. Urethral lappet with 2 forks. Dorsal papilla with single tip. Proximal baculum bony with a semicircle-shaped base.

Distal baculum also bony, short, base bugled largely. Lateral bacular processes cartilaginous and short (Supplementary Fig. S2e).

*Reproduction*.—In late October and November, no adult males orchidoptosis and no females pregnant. Data not available for other months.

*Habitat*.—This species inhabits moist mixed coniferous broad leaf forest at elevations about 3150 m a.s.l., tree height approximately 15-–18 m, 60% coverage. Humus 5–10 cm thick. Shrubs 3m and 30% coverage, grass 30 cm height and 15% coverage.

**Family Cricetidae Rochebrune, 1883**

**Subfamily Arvicolinae Miller, 1906**

**Genus *Neodon* Horsfield, 1841**

***Neodon bershulaensis* sp. nov.**

**Bershula Mountain vole**

*Holotype*.—Adult male, field number XZ10010 (Museum number: SAF11530), collected by Liao Rui on 3 Mach 2011. Specimen was preserved at the Sichuan Academy of Forestry as a skin, cleaned skull, penis and tissue. External and cranial measurements (in mm) as follows (abbreviations in Material and Methods): HBL 116.0 mm; TL 53.0 mm; HFL 18.0 mm; EL 13.0 mm; SGL 27.56mm; SBL 25.87mm; CBL 26.81 mm; ZB 15.89 mm; IOW 4.35 mm; MB 12.53 mm; SH 9.90 mm; ABL7.51 mm; LMxT 6.48 mm; LMbT 6.52 mm; LM 19.96; M-M 5.625 mm; and OLLI 9.10 mm. Body mass 40g. Photos of skull, dentition, and mandible in (Supplementary Fig. S1f).

*Type locality*.—Ridong village, Bershula Mountains, Chayu County, southeast of Xizang, China. 98.12407° E, 28.58392° N, elevation 3750 m a.s.l. Specimen was collected with a steel trap (Jiangxi Mouse Devices Factory) under fallen fir deadwood, with thick humus.

*Paratypes*.—3 specimens (2males, 1 females); 1 intact adult, field number: CHYRD-03♀; two intact juveniles, field numbers: CHYRD-04♂, CHYRD-02-001♂.

*Distribution*.—Known from the type locality only, Ridong village, Chayu County, southeast of Xizang.

*Etymology*.—Species epithet for famous Bershula Mountains, where type locality, Ridong is at its foot. Bershula Mountains is a famous mountains and with high diversities. The name highlights the importance of conserving this area for its endemic biodiversity.

*Diagnosis*.—Medium body, average length 108 mm (adult); hind feet 18–20 mm (average 19 mm). Tail length averages 53.5 mm, 49.5% of HBL. First lower molar with 5 closed triangles, 6 inner and 4 outer angles. 1$^{st}$ upper molar with 4 inner and 3 outer angles in 70% specimens; other 30% with 3 inner and 3 outer angles. 2$^{nd}$ upper molar with 3 inner and 3 outer angles. 3$^{rd}$ upper molar with 4 inner and 3 outer angles (Supplementary Fig. S1f).

*Description*.—Pelage from head to hip uniform grey-brown. Entire back covered with fine, dense, velvet hair. Ventral hairs grey-white, blushing with brown tinge. Transition between dorsal and ventral pelage vague. Ears project above pelage slightly, covered with short grey-brown hairs. Tail bicolor; dorsum grey-black, ventral grey-white; dorsal hairs slightly longer. Dorsal surface of forefoot and hindfoot grey-black. Claws grey-white, upper surface of nail grey-black. Five palmar and 6 plantar pads. Females with 1 pair of inguinal and pectoral mammae.

Skull relatively sturdy, dorsal profile straight, brain case flattened (Supplementary Fig. S1f). Nasal short, broad anteriorly narrowing posteriorly. Parietal irregular with lateral protrusion. Interparietal broad, irregular rectangle, mid-anterior part protruding forward. Interorbital and temporal ridges absent. Zygomatic arches slender, middle part slightly broader. Auditory bullae moderately sized. Incisory foramen relatively long and broad. Posterior palate typical of *Microtus*, with 2 obvious lateral pits. Many mini-foramen in palate and pterygoid. Mandibles medium (Supplementary Fig. S1f).

Upper incisors orange. Molars rootless. $1^{st}$ upper molar of holotype with 4 closed triangles after anterior transverse space, 4 outer and 3 inner angles, but inner fourth vestigial; 30% specimens with 3 inner an 3 outer angles. $2^{nd}$ upper molar with 3 closed triangles after the anterior transverse space, forming 3 inner and 3 outer angles. $3^{rd}$ upper molar with 4 inner and 3 outer angles (Supplementary Fig. S1f6). $1^{st}$ lower molar with 5 closed triangles and a trefoil anterior tooth cap, this tooth with 6 inner and 4 outer angles. $2^{nd}$ and $3^{rd}$ lower molars with 3 outer and 3 inner angles (Supplementary Fig. S1f7).

Glans penis (Fig. 4, Supplementary Fig. S2f) slender, pole-like and with ventral groove. Outer crater with 1 obvious papilla on both sides. Urethral lappet with 2 forks. Dorsal papilla with single tip. Proximal baculum bony with a semicircle-shaped base. Distal baculum cartilaginous and tongue-shaped, tip pointed. Lateral bacular processes also cartilaginous.

*Reproduction*.—In mid-October and late March, no adult males orchidoptosis and no females pregnant. Other months without reproduction information.

*Habitat*.—This species inhabits shrubs with sparse firs at elevations of 3450–3750 m a.s.l., tree height approximately 15 m, and 15% coverage. Shrubs 2–3m and with 40% coverage. Understory, humus 10–15cm thick and grass 20–50cm, 20% coverage.

#### SUPPLEMENTARY FIGURES CAPTIONS

**Supplementary Figure S1** Skull comparisons for the unidentified taxa. 1. Ventral view, 2. Dorsal view, 3. Lateral view, 4. Ventral lower jaw, 5. Lateral lower jaw, 6. Upper toothrow and 7. Lower toothrow of six unidentified species of *Neodon*. A. unidentified taxon from north of Yarlung Zangbo River. B. unidentified taxon distributing between south of Yarlung Zangbo River and north of Namchabarwa moiuntains. C. unidentified taxon from southern Namchabarwa Mountains. D. unidentified taxon from Chayu County. E. unidentified taxon from Chayu County. F. unidentified taxon from Bershula mountains.

**Supplementary Figure S2** Comparison of the glans penis of eight voles. A. *N. sikimensis*; B. *N. namchabarwaensis*; C. *N. liaoruii*; D. *N. chayuensis*; E. *N. bomiensis*; F. *N. bershulaensis*; G. *N. forristi*; H. *N. shergylaensis*. Numbered views are 1. Glans; 2. Midventral cut view; 3. Urethral lappet; 4. Dorsal papilla.

**Supplementary Figure S**3 Heatmap of nuclear gene coverage

**Supplementary Figure S4** Nuclear K2P distances between *Neodon* species. The average genetic distances of all available nuclear genes between each pair species were used to draw this heatmap. The species were arranged into three groups (clade1, clade2 and clade3. refer to Fig. 5 for clade information). Bold text: new species described herein.

**Supplementary Figure S5** Phylogenetic trees inferenced by *cox1* and *cytb*.

**Supplementary Figure S6** Mitochondrial trees and species delimitation. Mitochondrial phylogenetic tree from a. RAxML b. Species delimitation results, and c. MrBayes tree.

**Supplementary Figure S7** Nuclear tree and species delimitation. Phylogenomic tree obtained using ASTRAL-III method and branch lengths were re-estimated in units of substitutions per site using ExaML. The topology was identical using both coalescent-based methods (ASTRAL and SVDquartets).

**Supplementary Figure S8** Phylogenetic trees inferenced using nuclear genes from two concatenated methods. a. RAxML and b MrBayes.

**Supplementary Figure S9** Distributing pattern of three clades of Neodon.

**Supplementary Figure S10** dN and dS trees. a. Mitochondrial tree inferenced from PAML with branch model. b. Distribution of Mitochondrial dN, dS and dN/dS. c. Nuclear tree inferenced from PAML using 100 nuclear genes with branch model. d. Distribution of nuclear dN, dS and dN/dS.

**Supplementary Figure S11** ILS results. a. Species tree with branch numbers. b. Relative frequency. Main topologies are shown in red, and the other two alternative topologies are shown in blue. Dotted lines indicate the 1/3 threshold. Title of each subfigure indicates the label of the corresponding branch on the tree on the right. c. Relationship between ILS occurrence frequency (ln value) and inner-node branch length of time tree.

#### SUPPLEMENTARY TABLES CAPTIONS

**Supplementary Table S1** Information of samples

**Supplementary Table S2** Measurements of morphological characteristics

**Supplementary Table S3** Calibration points

**Supplementary Table S4** Information of high coverage gene set

**Supplementary Table S5** Intra-species distances of Mitochondrial 13 PCGs

**Supplementary Table S6** Congenic inter-species distances of Mitochondrial 13 PCG

**Supplementary Table S7** Inter-genera distances of Mitochondrial 13 PCGs

**Supplementary Table S9** dN and dS ratio of mitochondrial genes estimates from PAML with branch model

**Supplementary Table S10** dN and dS ratio of nuclear genes estimates from PAML with branch model

**Supplementary Table S11** Fit for DEC and DEC+j models of ancestral range estimates

#### LIST OF SUPPLEMENTARY APPENDICES

**Supplementary Figures** Supplementary Figures S1-S11

**Supplementary Tables** Supplementary Tables S1-S11

**Supplementary Appendix S1** Supporting Online Material

**Supplementary Appendix S2** Dataset 1 with mitochondrial genes

**Supplementary Appendix S3** Dataset 2 with all nuclear genes

**Supplementary Appendix S4** Dataset 3 with 100 nuclear genes

**Supplementary Appendix S5** Dataset 4 with phase 1 sites of all nuclear genes
