## Supplementary figures and images for "Out of the Qinghai-Tibetan Plateau and get flourishing - the evolution of *Neodon* voles (Rodentia: Cricetidae) revealed by systematic sampling and low coverage whole genome sequencing"

### Supplementary Fig. S1

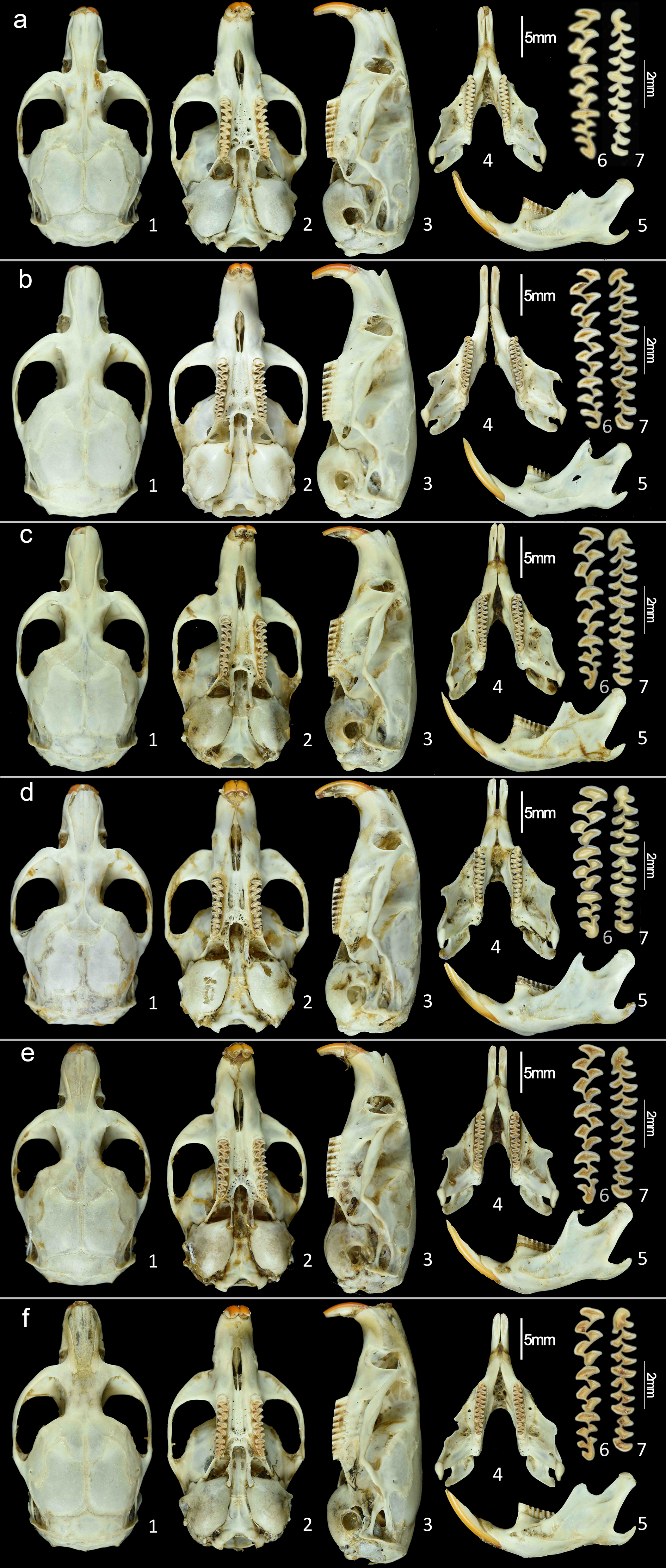

### Supplementary Fig. S2

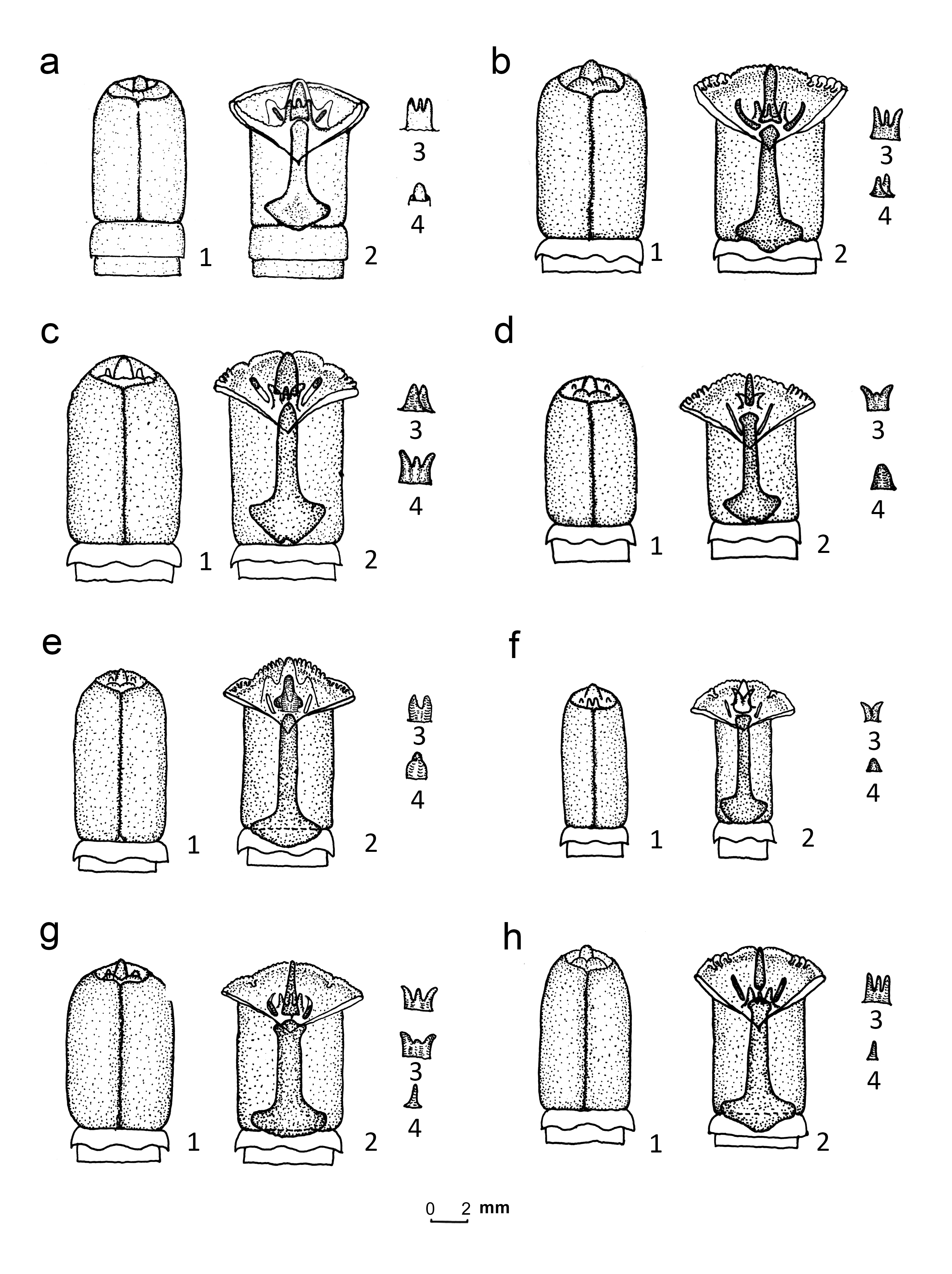

### Supplementary Fig. S3

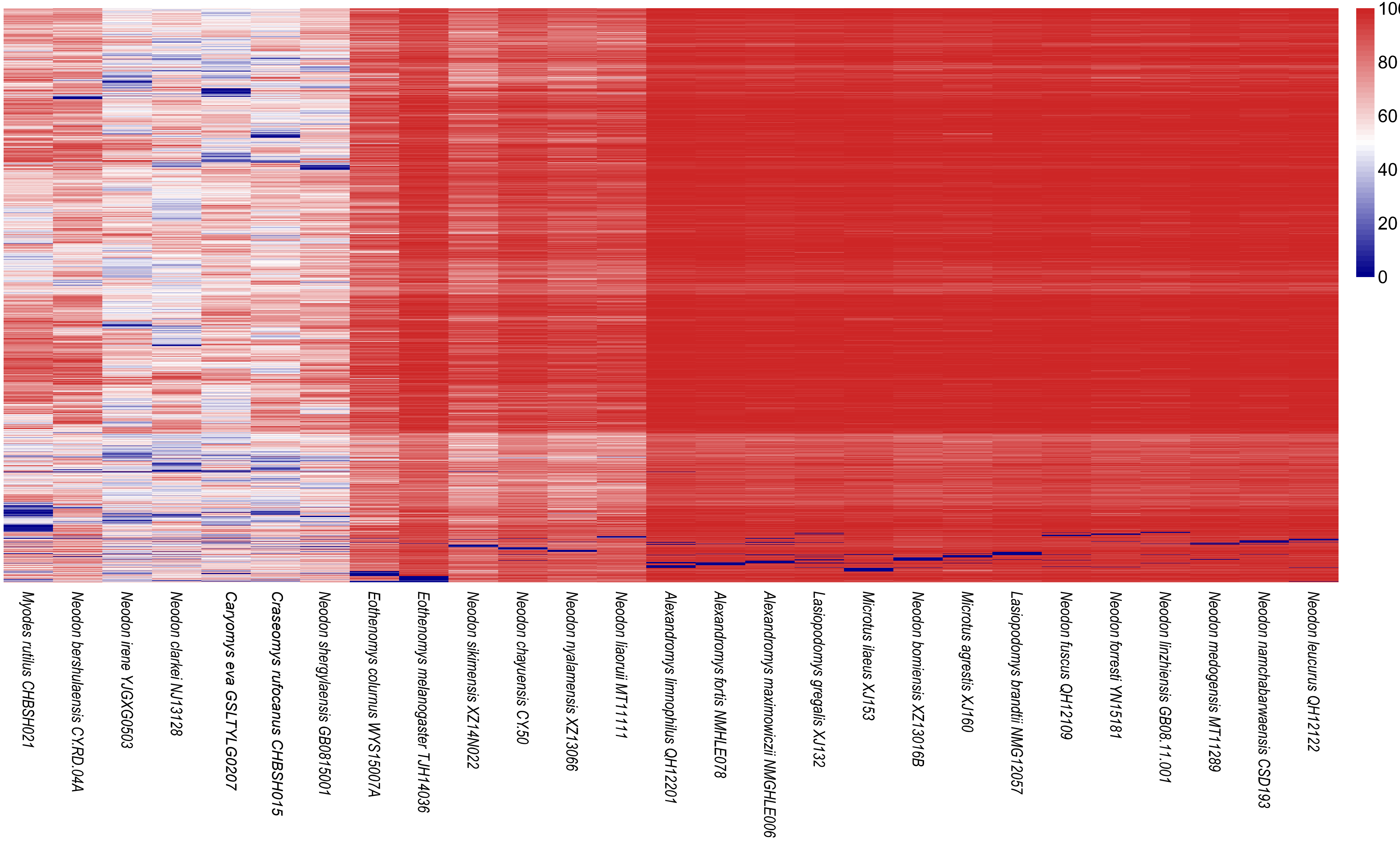

### Supplementary Fig. S4

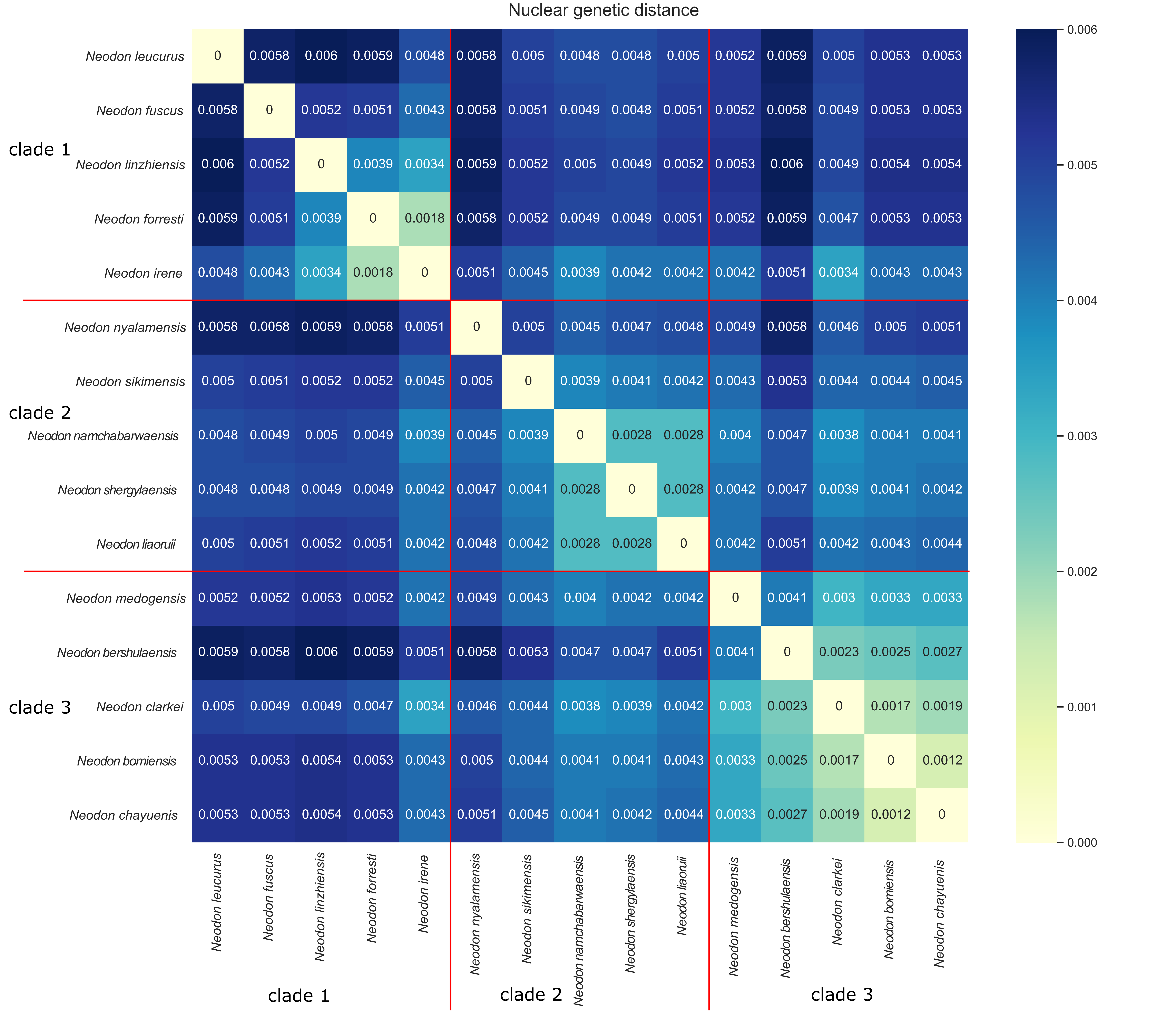

### Supplementary Fig. S5

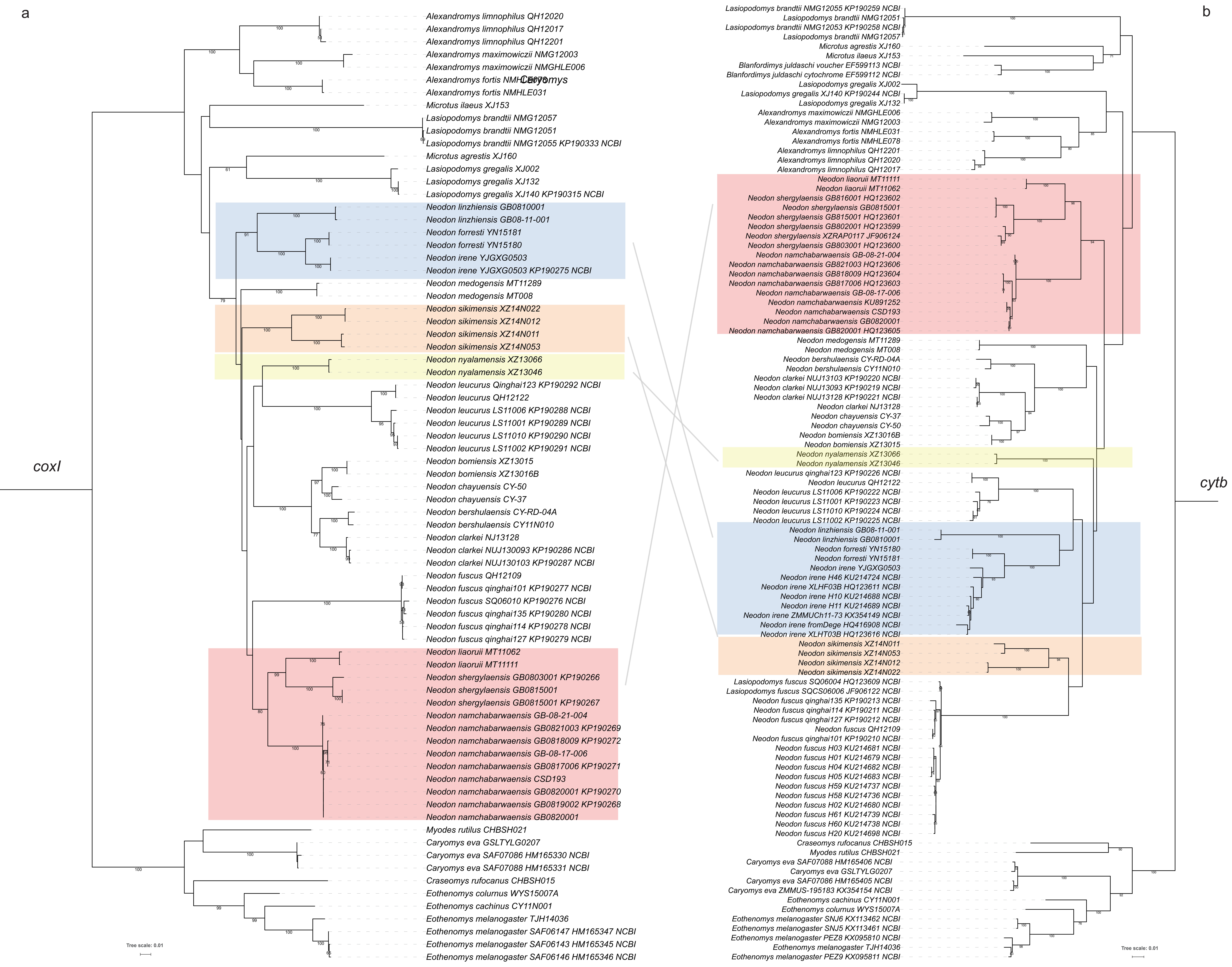

### Supplementary Fig. S6

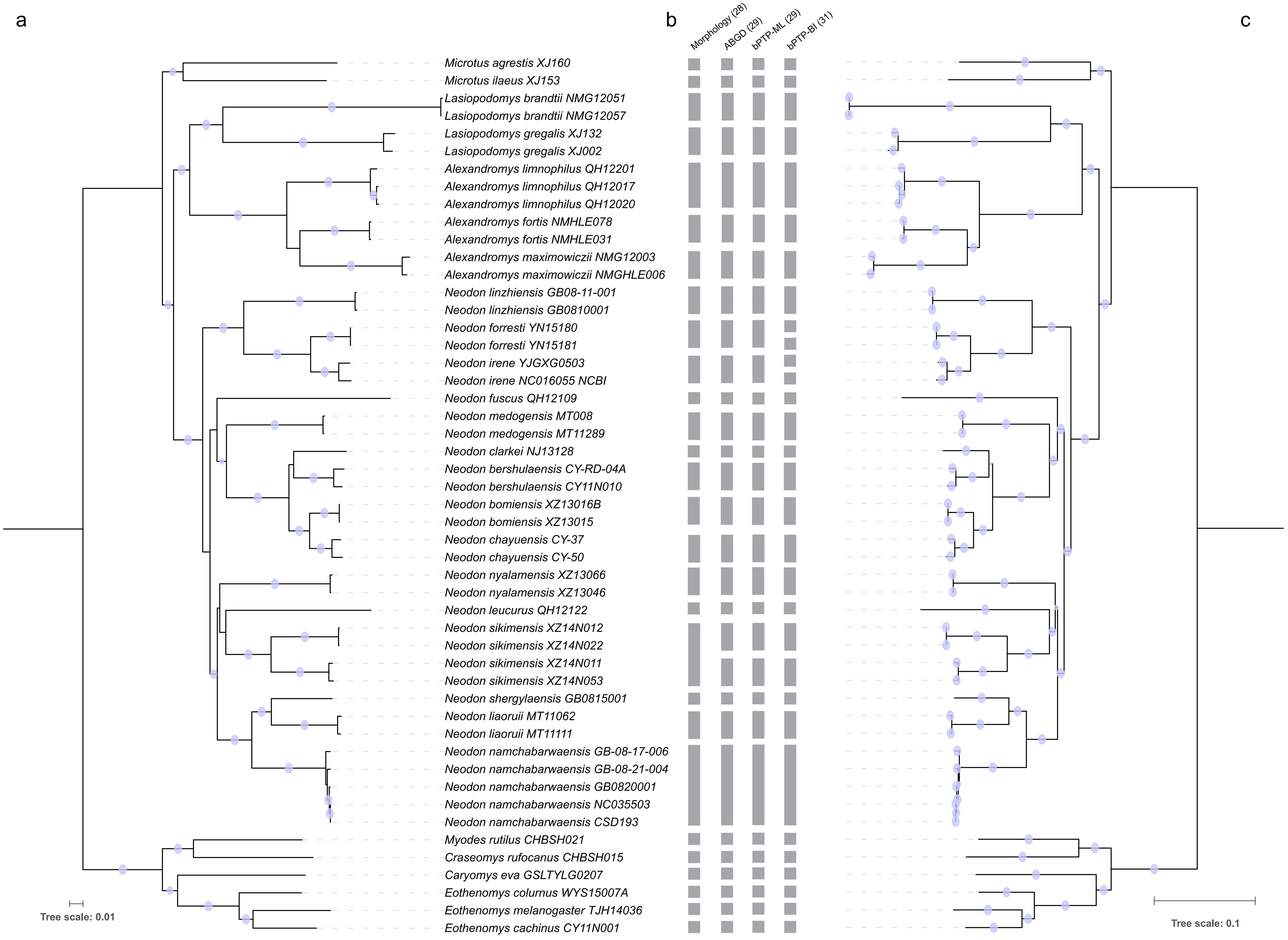

### Supplementary Fig. S7

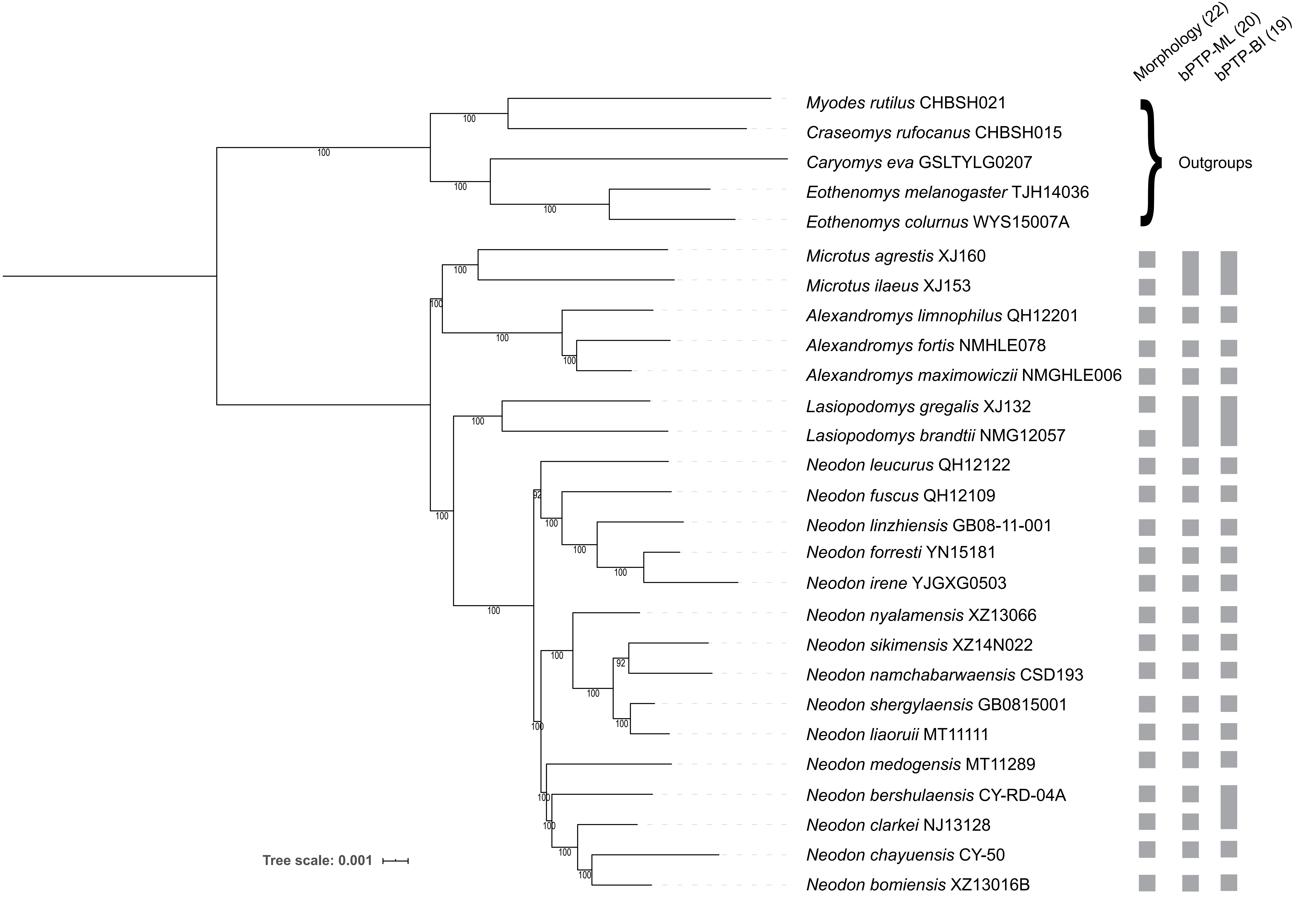

### Supplementary Fig. S8

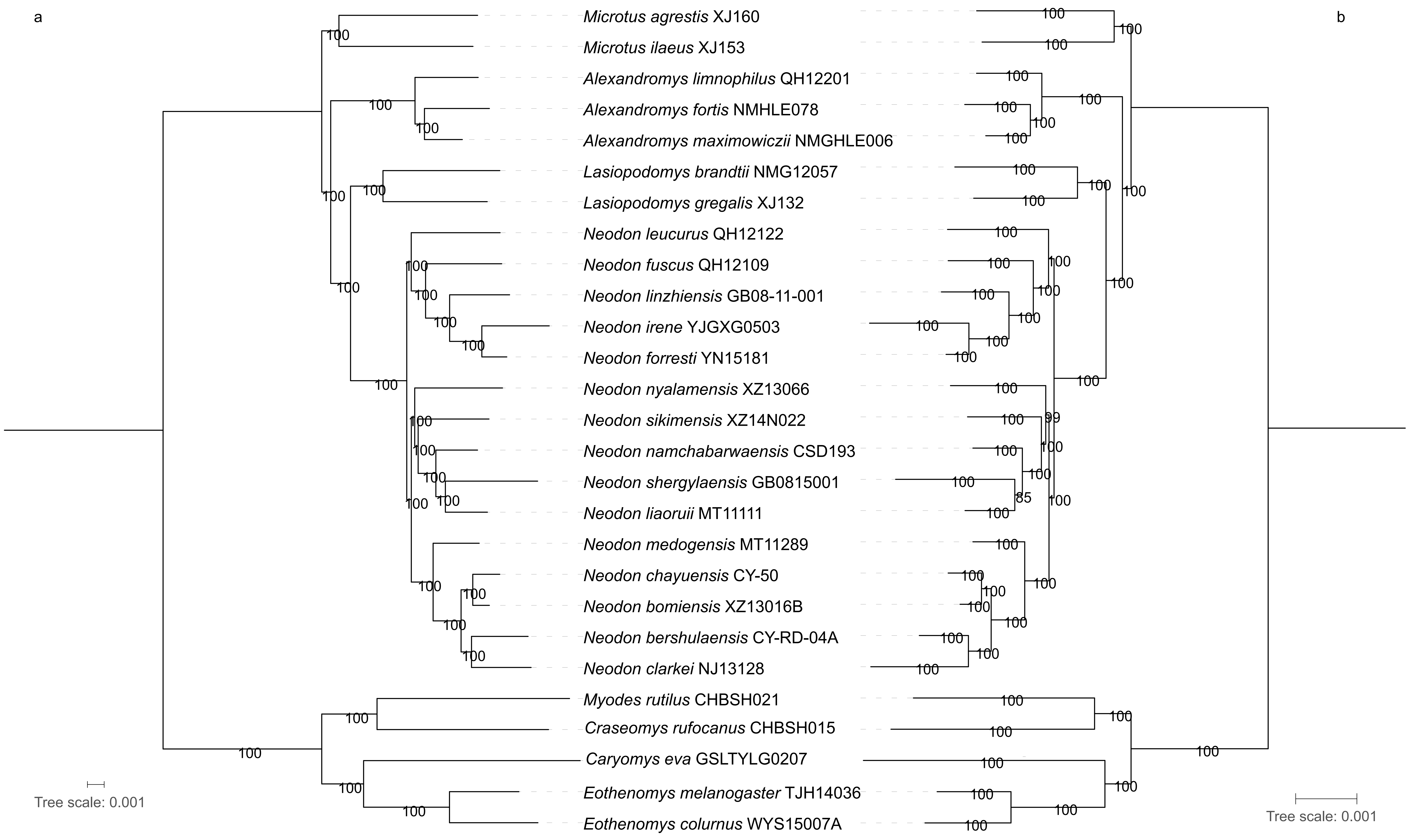

### Supplementary Fig. S9

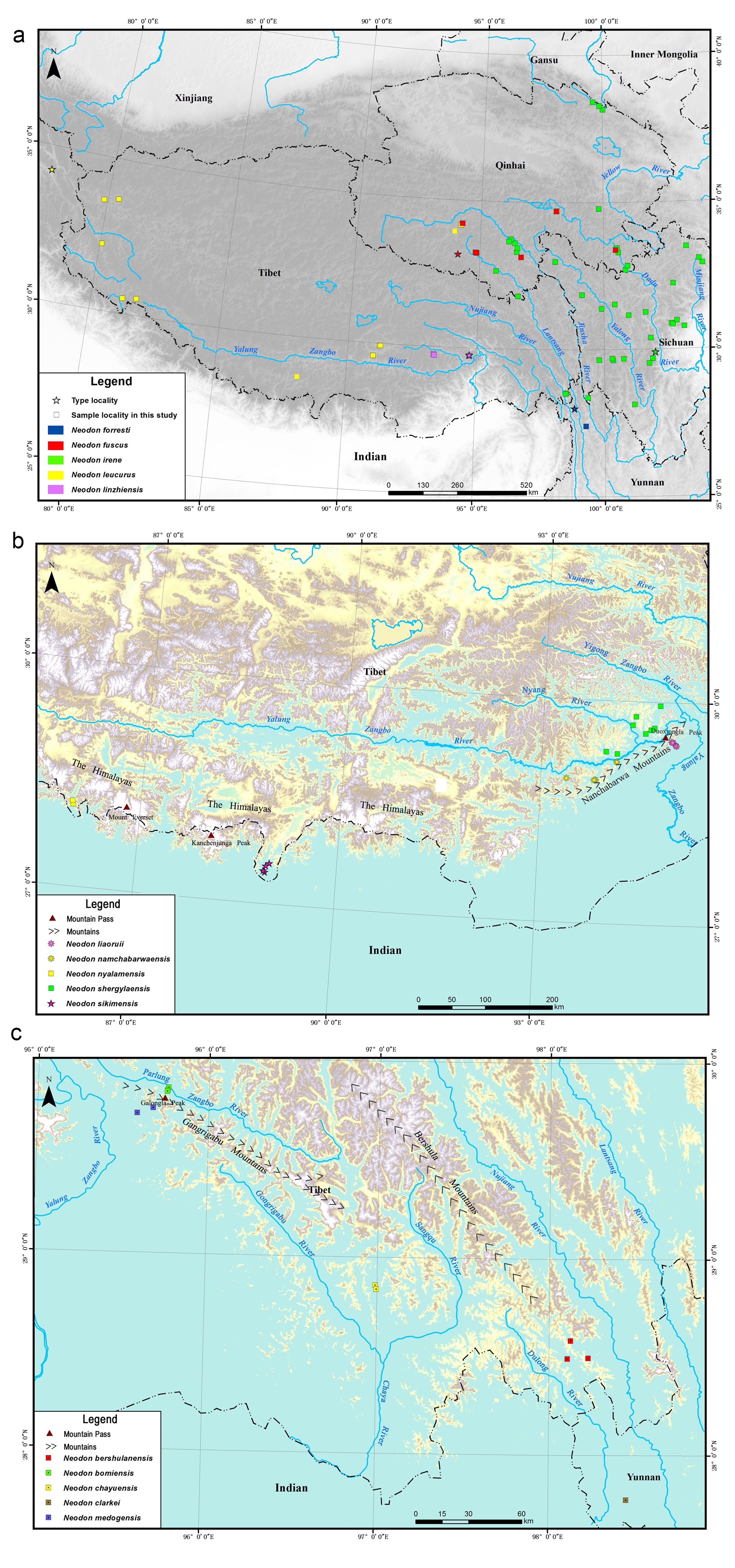

### Supplementary Fig. S10

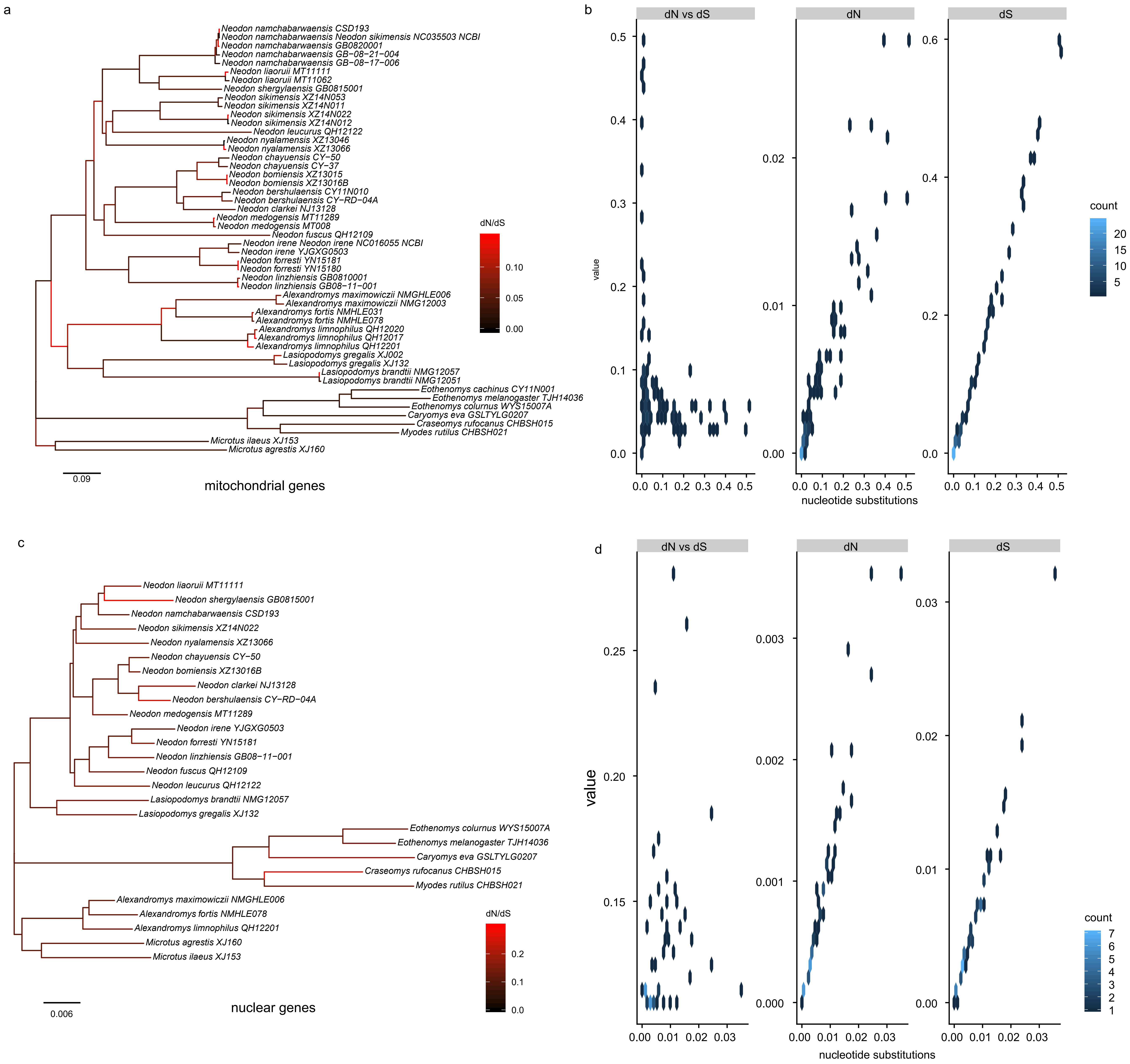

### Supplementary Fig. S11

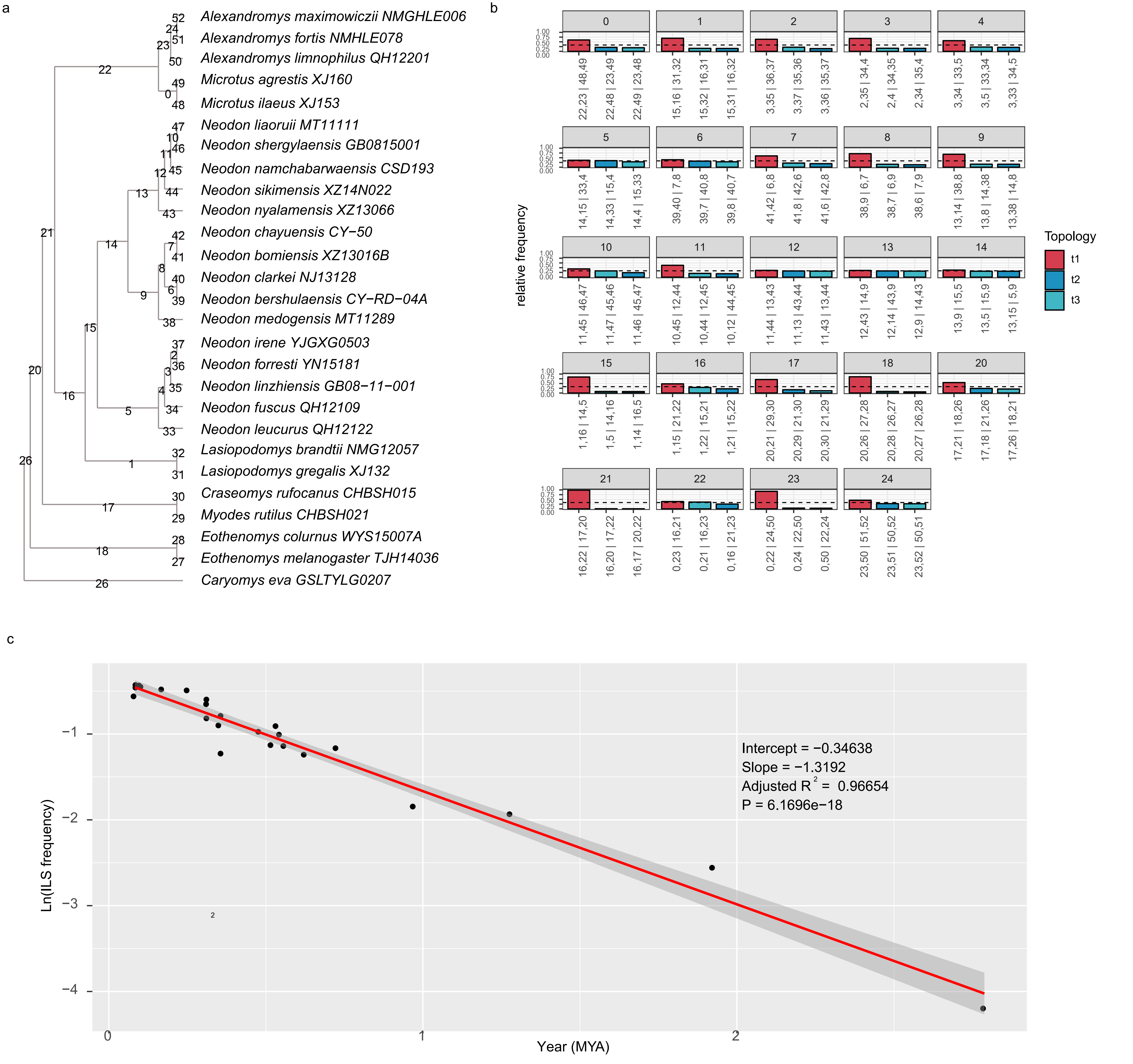
